# Size control and oscillations of active droplets in synthetic cells

**DOI:** 10.1101/2024.07.24.604607

**Authors:** Judit Sastre, Advait Thatte, Alexander M. Bergmann, Michele Stasi, Marta Tena-Solsona, Christoph A. Weber, Job Boekhoven

## Abstract

Oscillations in the formation and dissolution of molecular assemblies inside living cells are pivotal in orchestrating various cellular functions and processes. However, designing such rhythmic patterns in synthetic cells remains a challenge. Here, we demonstrate the spontaneous emergence of spatio-temporal oscillations in the number of droplets, size, and position within a synthetic cell. The coacervate-based droplets in these synthetic cells sediment and fuse at the cell’s bottom. Through a newly discovered size control mechanism, the sedimented, large droplets shrink by expelling droplet material. The expelled molecules nucleate new droplets at the top of the synthetic cell, which grow and sediment again. These oscillations are sustained by converting chemical fuel into waste and can continue for hundreds of periods without evidence of fatigue. Strikingly, the period of the oscillation is in the minute’s regime and tunable. The design of oscillating organelles in synthetic cells brings us closer to creating more life-like materials and *de novo* life.

---

Oscillations in cells are essential for precisely regulating cellular and physiological processes in space and time.^1^ Oscillations are not limited to fluctuation in concentrations of biomolecules but can also involve entire biomolecular assemblies.^2^ An example is the Min oscillatory system in E. coli cells that regulates cell division using the Min-protein complex that oscillates from pole to pole.^3^ Systems chemistry aims to mimic and engineer systems with the sophistication and complexity of biology in synthetic systems.^4,5^ Engineered oscillations in synthetic cells could function as autonomous time-keeping systems that regulate the internal organizations of the synthetic cell. Nevertheless, designing oscillations in synthetic systems remains challenging. Several oscillators exist in which the concentration of components oscillate, so-called chemical oscillators. Classical, well-established oscillators like the Belousov-Zhabotinsky-reaction^6^ and others^7^ are based on inorganic compounds, making them incompatible with synthetic cells. Oscillators based on organic compounds can be designed using enzymatic reaction networks^8^ and completely non-biological counterparts.^9^ Self-assembling molecules can be coupled to these oscillating chemical reactions, like a pH-clock reaction,^10^ to produce oscillating assemblies like micelles,^11^ vesicles,^12^ or hydrogels.^13^ However, oscillators in which the properties and function of molecular assemblies are direct components of the oscillating reactions, as observed in biology, remain sparse.^14,15,16^

Here, we developed synthetic cells endowed with active droplets. When the cells are constantly supplied with chemical energy, they produce active droplets that spontaneously oscillate in their organelle number, size, and position with a tunable period in the minute range. We found that the oscillations rely on a combination of a recently proposed size control mechanism,^17,18^ and gravitationally induced fusion of droplets. Size control means these active droplets tend to evolve to an optimal size, making droplets grow when too small and actively expelling material when too large. When we inhibit droplet fusion, the average size and number of droplets are controlled, contrasting droplets in equilibrium, which would settle for one droplet per synthetic cell. In contrast, when fusion is possible, the fused, large droplets shrink at the bottom of the cell, which induces the nucleation of new droplets at the top of the synthetic cell, which grow and sediment again. The period of the oscillation is in the minute’s regime and tunable. These oscillations are sustained by converting chemical fuel into waste and can continue for hundreds of periods without evidence of fatigue.

**Scheme 1.**
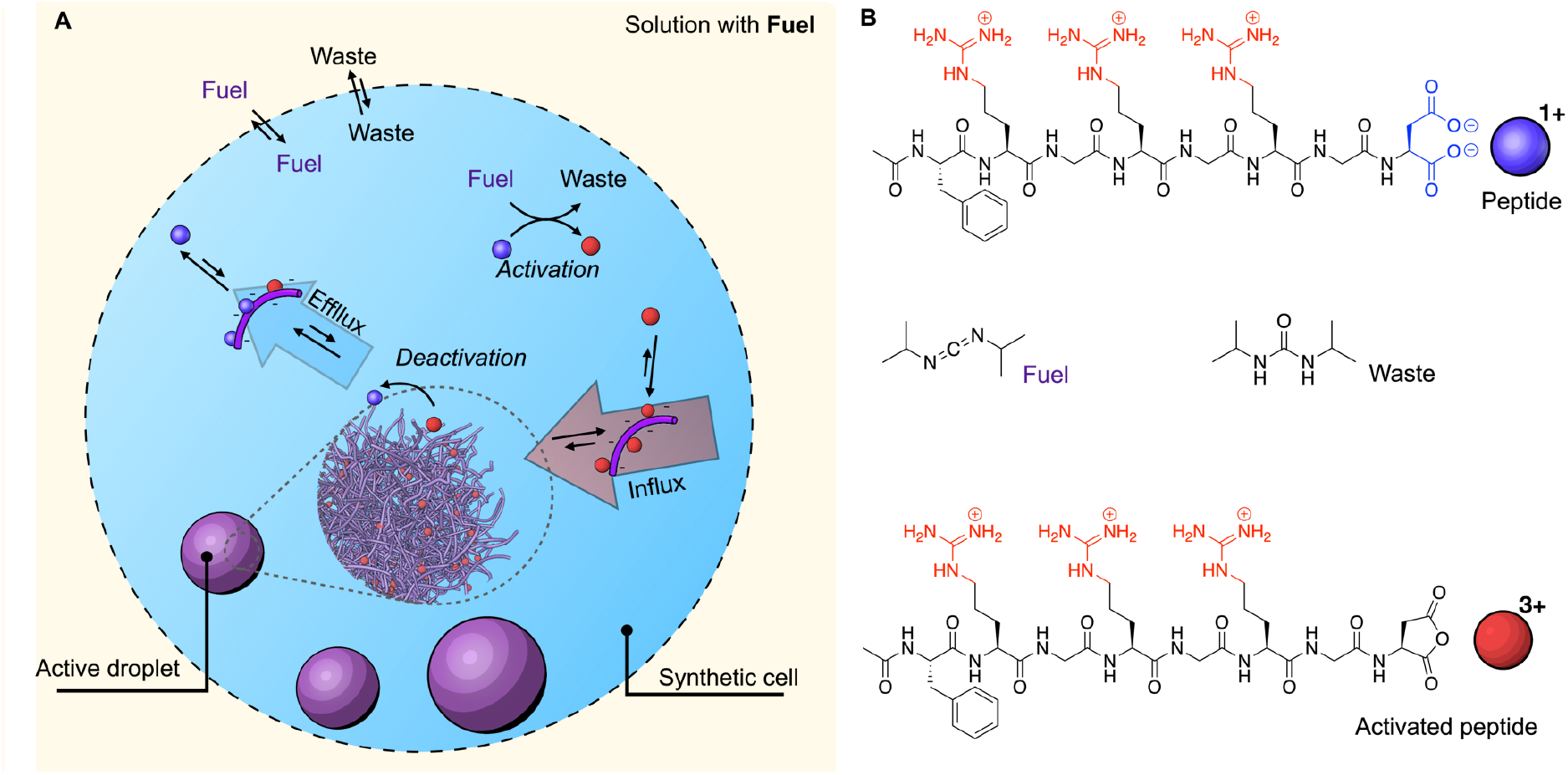
Active droplets are maintained out of equilibrium in a synthetic cell. A) The experimental setup used to sustain droplets in a steady state. An aqueous droplet (synthetic cell) with peptide and polyanion is created in a liquid of perfluorinated oil with fuel. Fuel exchange at the oil-water interface leads to a steady state that produces chemically fueled droplets. **B**) The chemical formulas of the reactants involved in the chemical reaction cycle.

## Oscillations in synthetic cells

The synthetic cells are formed by combining an aqueous solution with a peptide and a polyanion with a perfluorinated oil that contains a diisopropyl carbodiimide (DIC) as “fuel”.^19^ Mixing the two solutions creates water drops in the perfluorinated oil of 10-25 μm in radius as synthetic cells (Scheme 1A). At the interface of the synthetic cell, fuel exchanges with the oil, saturating the aqueous synthetic cell with fuel required to drive a chemical reaction cycle that produces active droplets,^20,21,22^ *i*.*e*., droplets made of transient droplet material.^23,24,25^ Specifically, the fuel continuously activates AcF(RG)_3_D-OH (peptide, Scheme 1B) and converts it into its corresponding cyclic anhydride (activated peptide, Scheme 1B). The activated peptide has a half-life of roughly a minute before it spontaneously deactivates through hydrolysis and reverts to its initial peptide state. The peptide is designed to be well-soluble in the synthetic cell. However, in its activated state, it is cationic with an overall charge of 3+ and thus can form a complex coacervate with a polyanion (17 kDa polystyrene sulfonate, PSS). Consequently, the synthetic cells continuously produce transiently active material for active droplets at the expense of fuel that exchanges at the cell’s interface. In previous work, we established that the peptide and fuel reside mostly in the aqueous phase outside these droplets, and thus, droplet material activation occurs predominantly outside.<u>^19^</u> The activated peptide partitions mostly inside the droplets, and thus, deactivation occurs predominantly inside the droplets. This asymmetry of activation and deactivation leads to an influx of activated droplet material and an efflux of deactivated droplet material over the interface. These fluxes are the prerequisite for the occurrence of oscillations.

We imaged the entire synthetic cell over time by obtaining Z-stacks in confocal microscopy (Figure 1A and B, Supplementary Movie 1). We were surprised to find that the droplets showed oscillatory behavior—the droplets nucleated at the top of a synthetic cell, grew to a radius of roughly 1 μm, sedimented, and, at the bottom of the reactor, fused to form a few large droplets. Then, after about a minute, the process repeated, *i*.*e*., new droplets nucleated and sedimented. We measured the projected surface area of each droplet, from which we calculated the radius of each droplet with time. We calculated the combined volume of all droplets in the synthetic cell from these radii, which was relatively stable, pointing towards a steady state in the amount of droplet material in the entire synthetic cell (Supplementary Figure 1). Then, we quantified the total droplet amount in the synthetic cell’s upper and lower half, which showed that oscillations in the lower and upper part are phase-shifted (Figure 1C)— when the total droplet volume in the top half peaked, the bottom half’s total droplet volume usually showed a valley and *vice versa*. Remarkably, the synthetic cells oscillated for hours without signs of fatigue (Figure 1D).

**Figure 1.**
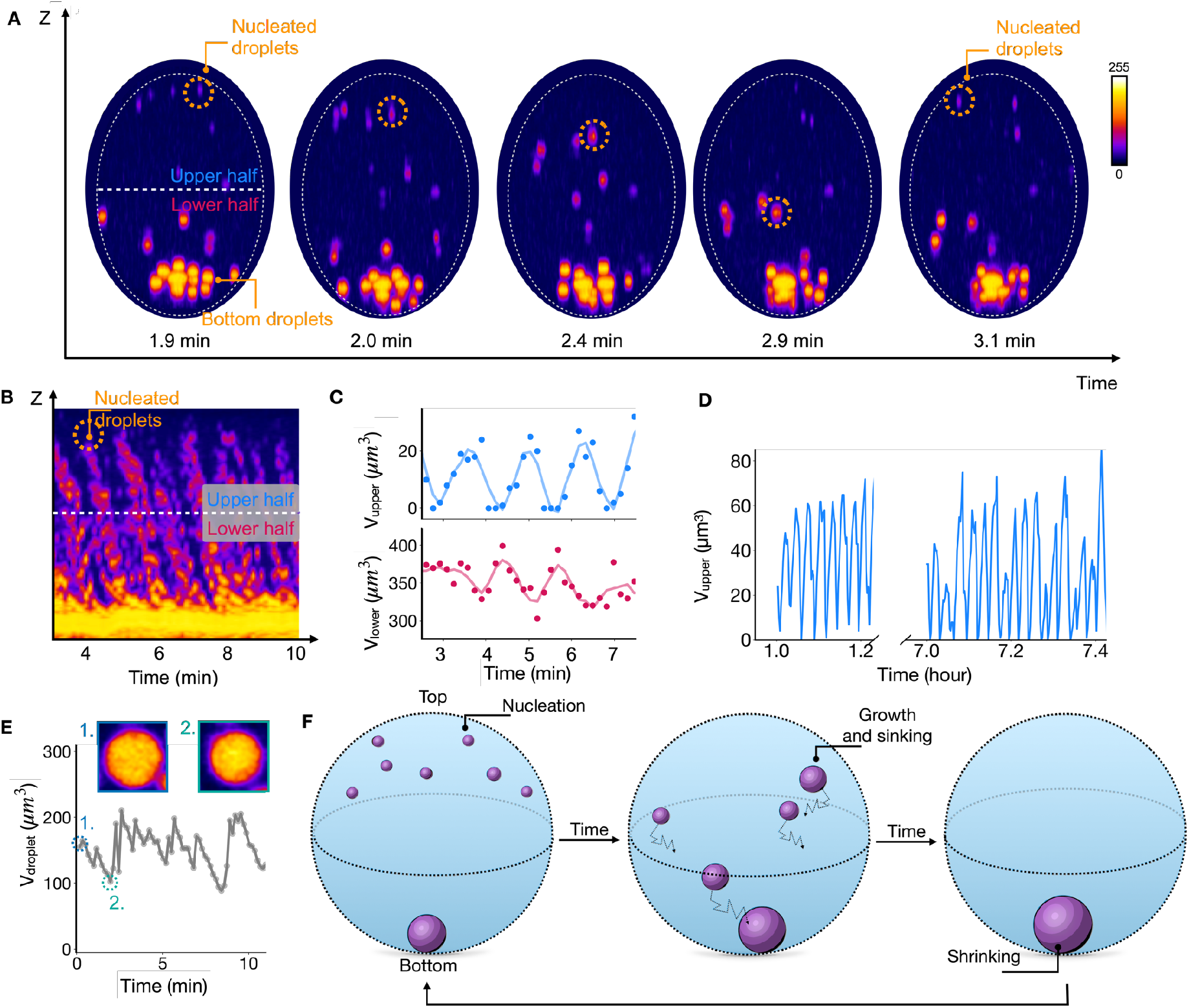
Oscillations in the position of organelles in synthetic cells. **A.** Sideviews of a synthetic cell prepared from confocal Z-stacks. The synthetic cell contains 10 mM peptide, 5 mM PSS polyanion, and 200 mM MES buffer and is embedded in oil with 250 mM DIC as fuel. This fuel is continuously exchanged at the droplet-oil interface. The number and the position of the organelles oscillate with time. **B**. Kymograph of the data from A. **C**. The volume of all organelles combined in the upper or lower half of the synthetic cell is described in A and B. The markers represent the measured data, and the line represents the moving average of five data points. **D**. The same as C, but the data is displayed for several hours. Data corresponds to a different cell size, hence the difference in values (Supplementary Figure 2) **E**. The radius of a selected droplet at the bottom of the synthetic cell. The two selected micrographs show the rapid shrinkage of the organelle. **F**. Schematic representation of the three-phase oscillations: 1) Nucleation, 2) growth and sedimentation, 3) shrinkage.

The initial part of an oscillation can be explained intuitively. Complex coacervate droplets have a low interfacial surface tension, facilitating fusion.^26^ Since their density is larger than their surrounding phase, droplets sediment.^27,28^ The next step is not intuitive, *i*.*e*., the large droplet(s) at the bottom have to shrink to provide material for the renucleation of droplets at the top. Indeed, when we tracked the volume of a large droplet at the bottom of the reactor, we observed rapid growth events through fusion followed by droplet shrinkage of the big droplet(s) at the bottom (Figure 1E). Such a behavior is thermodynamically unfavorable since the decrease in droplet size and the renucleation of small droplets elsewhere, while keeping the total droplet volume the same, increases the total droplet surface area and, thereby, the free energy of the system. From this observation, we conclude that the droplet shrinkage in the last step of the oscillation cycle must be active behavior.

## Mono-disperse active emulsion: Active droplets evolve to the same optimal size

We tested whether the counterintuitive droplet shrinkage was a system property or a result of the confined environment of the synthetic cell. Thus, we prepared a mm-thick layer of the peptide solution (and polyanion) and carefully placed the pure diisopropyl carbodiimide as fuel on top (Figure 2A). At the water-fuel interface, the fuel would exchange and saturate the aqueous buffer solution. We found the emergence of droplets throughout the thin water layer. The droplets fused rapidly and sedimented, which we quantified by analyzing the droplet number and radii (Figure 2B, Supplementary Figure 3 and Supplementary Movie 2). Excitingly, a closer look at the droplet radii revealed that, between fusion events, the radius of some droplets decreased, pointing to a combination of fusion and the droplet correcting its size by shrinking (Figure 2B). This result demonstrates that the shrinkage of large droplets is independent of the confinement of the synthetic cell.

**Figure 2.**
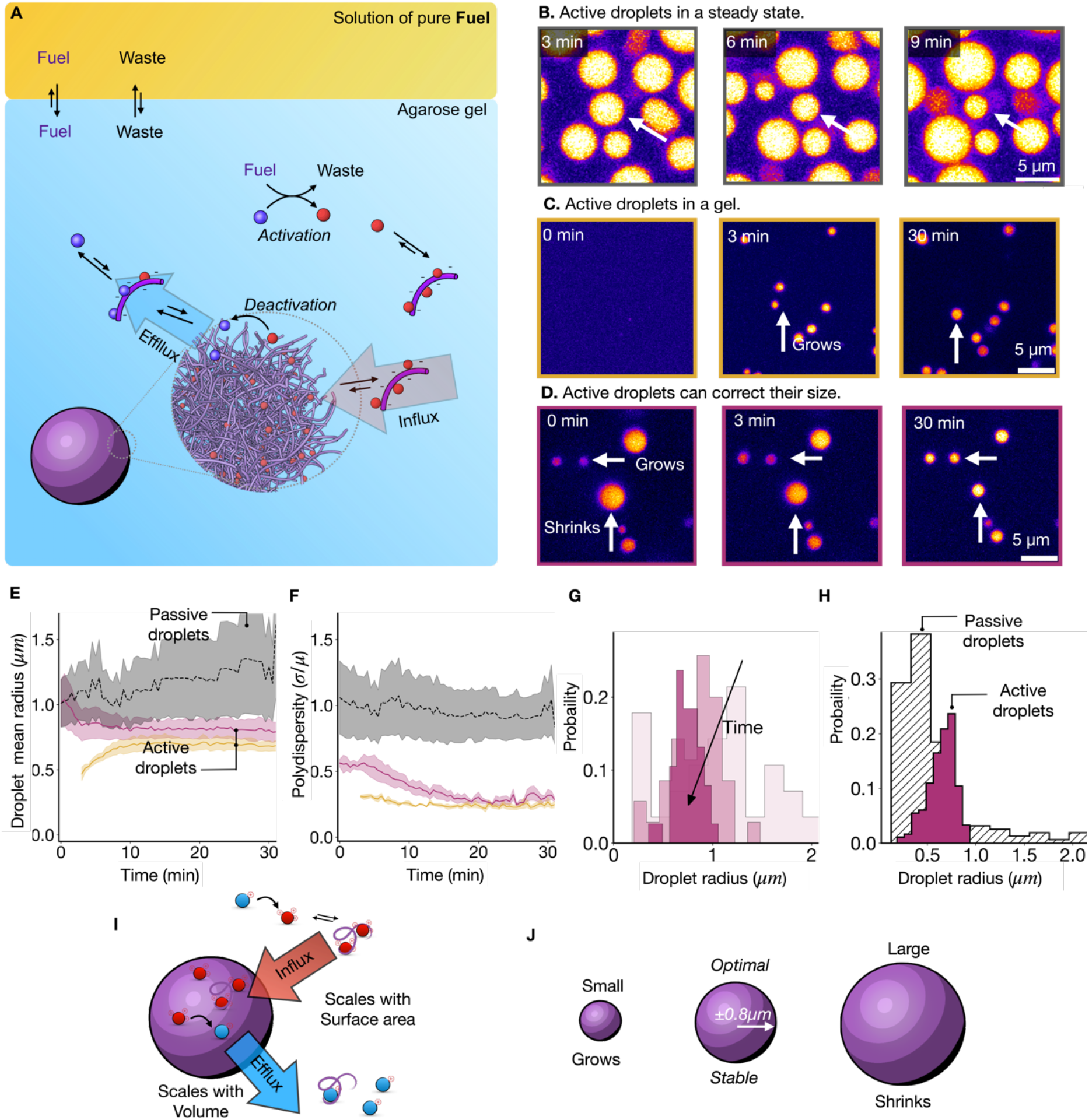
Active droplets control their size. **A**. Schematic representation of the experiment. **B**. Micrographs of active droplets are continuously supplied with chemical fuel. A solution of 14 mM peptide, 15 mM PSS, 0.2 μM sulforhodamine B in 200 mM MES buffer at a pH of 5.3 was covered with a layer of DIC. The droplets were imaged by confocal microscopy. **C**. Micrographs of active droplets under the same conditions in A, now embedded in a 0.5% agarose hydrogel to prevent fusion. **D**. Micrographs of active droplets under the same conditions in C, now spiked with 20 mM EDC to induce pre-nucleation and fusion of droplets. **E-F**. The average droplet radius (**E**) and polydispersity (**F**) for passive droplets (grey), active droplets trapped in a gel network (yellow), and pre-nucleated, active droplets trapped in a gel (purple). The active droplets evolve to the same stable radius, whereas the passive ones steadily grow. **G-H**. Histograms of the droplet radius distributions as a function of time for the experiment described in D (**G**) or for active and passive droplets in a gel after 60 minutes (**H**). **I**. Schematic representation of the size control mechanism. Droplet material is activated outside of the droplet, leading to a surface area-dependent influx. Droplet material is deactivated inside of a droplet, which scales with its volume, leading to a volume-dependent efflux. **J**. A Surface area-dependent influx and volume-dependent efflux imply an optimal droplet radius. Small droplets will grow, and large droplets will shrink towards that radius.

We reduced the complexity of the experiment by suppressing sedimentation and fusion—we performed the same experiments with 0.5% w/v agarose gel to suppress droplet fusion. After the fuel was carefully pipetted on top of the gel, droplets emerged, but no fusion was observed while the droplet number remained constant (Figure 2C, Supplementary Figure 4 and Supplementary Movie 3). Surprisingly, we tracked the radii of hundreds of droplets, which initially grew until they all stabilized at roughly 0.8 μm within 15 minutes (Figure 2E and Supplementary Figure 4A). We quantified the deviation of the mean radius to calculate the polydispersity index as a function of time (PDI = σ/μ, where σ is the standard deviation of the mean radius and μ is the mean radius), which settled at only 0.2 confirming a narrow radius distribution (Figure 2F). Thus, the active droplets withstand fusion because of the gel network. Most importantly, they also seem to withstand growth or shrinkage through Ostwald ripening, settling to a common optimal radius. We verified that the optimal radius of droplets was independent of the agarose concentration to ensure the growth suppression is not a result of the gel’s elasticity or mesh size (Supplementary Figures 5-7).^29^

The droplets seem to grow to one optimal radius. To scrutinize this, we tested whether large droplets could also shrink to the same optimal size by initiating an experiment with a wide initial droplet size distribution, i.e., whether the emulsion could correct the size of droplets. We prepared active droplets using the water-soluble fuel 1-ethyl-3-(3-dimethylaminopropyl)carbodiimide (EDC) before the droplets were trapped in the gel network (0.5% w/v agarose gel), allowing for fusion. Once the gel had been set, we again carefully pipetted diisopropyl carbodiimide on the gel as fuel to sustain the pre-nucleated droplets (Figure 2D, Supplementary Movie 4). Tracking the droplet radii unambiguously showed the size control mechanism at work—we found droplets with a radius greater than 0.9 μm shrunk while droplets with a smaller radius grew (Supplementary Figure 8). Moreover, the average droplet size decreased to a stable size of 0.9 μm, and the initially wide size distribution with a polydispersity index of 0.45 dropped within 20 minutes (Figure 2E and F). Notably, the experiments with or without pre-nucleated droplets converged to roughly the same radius and polydispersity (Figure 2E, yellow and purple). Histograms of the droplet’s radii distribution further corroborated the size control mechanism by gradually narrowing with time (Figure 2G). As a final control, we prepared passive droplets, *i*.*e*., a mimic of our active droplets but not regulated by chemical reactions (See Materials and Methods). We prepared the droplets in a gel matrix, tracked their radii, and found they increased with time (Figure 2E grey and Supplementary Figure 9). We observed that the polydispersity index was higher than for active droplets (Figure 2F grey). Besides, their radii increased, demonstrating that passive droplets cannot control their size and slowly grow, likely through Ostwald ripening (Figure 2H).

## The mechanism of size control

We propose that the size control mechanism is related to a previously proposed mechanism using theory that was never demonstrated experimentally to the best of our knowledge.<u>^17^</u> When activation occurs predominantly outside the droplets, the droplet material’s influx of activated droplet material scales with the droplet’s surface area (Figure 2I). When deactivation occurs predominantly inside the droplet, the efflux of deactivated droplet material scales with its volume. This interplay between surface area-dependent influx and volume-dependent efflux implies that large droplets (with a small surface area to volume ratio) will lose more deactivated droplet material than activated droplet material they take up—they shrink. In contrast, small droplets (with a large surface area to volume ratio) grow because they take up more material than they lose. Such a mechanism explains why all droplets converge to one optimal radius and why large droplets that have fused (e.g., in the oscillations) shrink.

To validate this mechanism, we used the previously determined chemical reaction cycle’s kinetics (Supplementary Table 1, Supplementary Figure 10) and the partitioning coefficient of the peptide, polyanion, and fuel as well as the diffusion constant of the peptides in the droplet phase and estimated the diffusion constant of the fuel and the peptides in solution<u>.^19^</u> We developed a theoretical framework with all these parameters that simulate droplets’ nucleation, influx, and efflux in a steady state like in our synthetic cells (Supplementary Method 2 and Supporting Table 2). We initiated a simulation in which droplets could grow under concentrations of fuel and precursor like in the experiments. Remarkably, the simulation showed that once a steady state was reached, the droplets reached a optimal droplet size, similar to the experimentally obtained values (Supplementary Figure 11). Moreover, we redistributed the droplet material amongst those droplets in the final state of the simulation to create a perturbed emulsion—some droplets were larger, and others were smaller than the optimal radius (Figure 3A). When we continued the simulation, the large droplets shrunk while small ones grew confirming the theoretically proposed size control mechanism.

**Figure 3.**
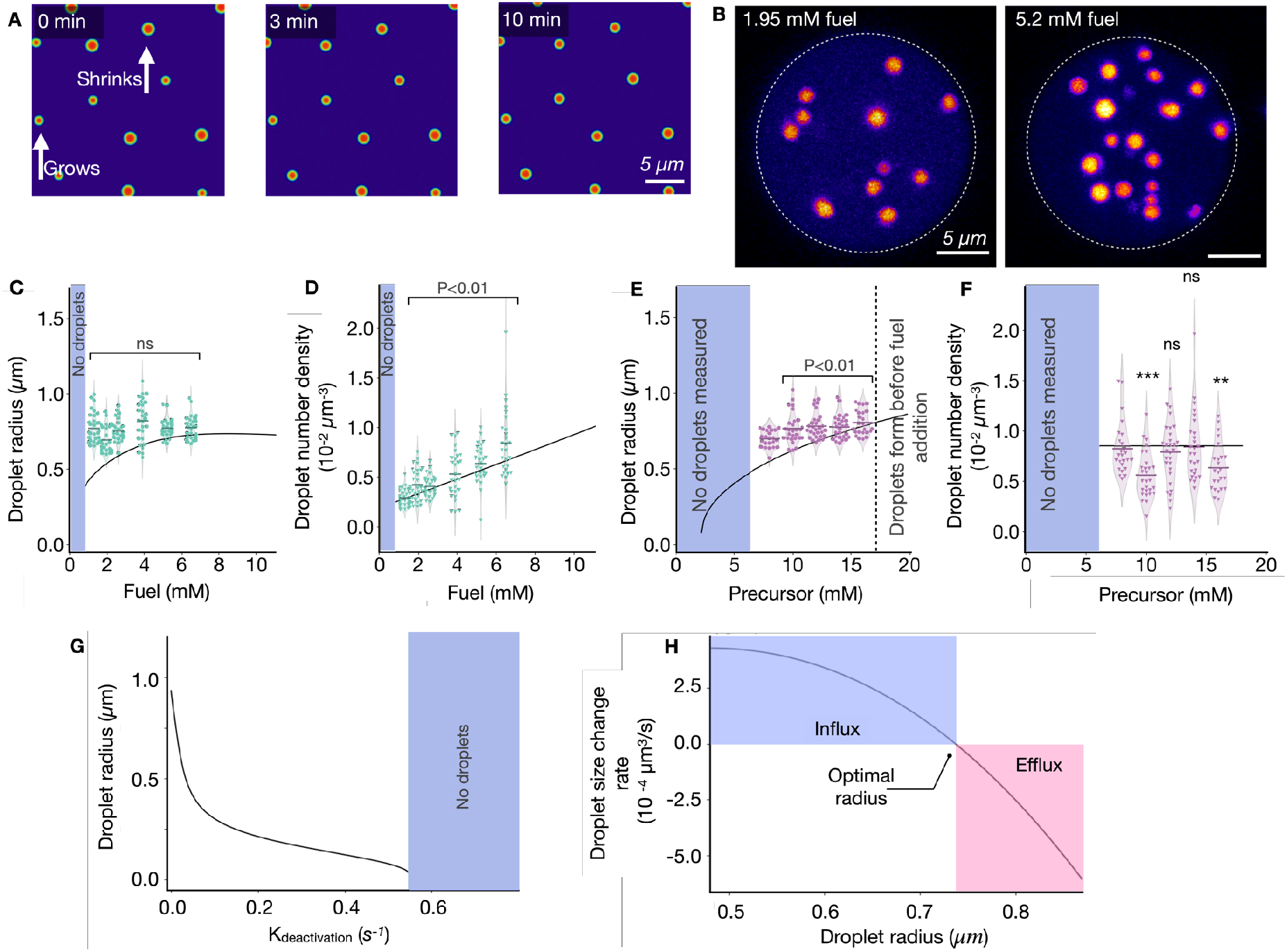
Parameters that regulate the optimal droplet radii and droplet density. **A**. Snapshots of the simulations of the kinetic model for active droplets (details in Supplementary Method 3) approaching their steady state after a size perturbation. When perturbing the steady state by increasing/decreasing randomly the size of droplets (left), decreased droplets grow, and increased droplets shrink toward their optimal radius (middle to right snapshot). **B**. Z-projections of synthetic cells with an agarose-based cytoskeleton for two different fuel concentrations (1.95 mM left, and 5.2 mM right) indicate that more active droplets form when increasing fuel. The conditions are 14 mM peptide, 15 mM PSS, 0.2 μM sulforhodamine B in 200 mM MES buffer at a pH of 5.3. **C-F**. The average droplet radius (C and E) and droplet number density (D and F) against the steady-state concentration of fuel (C, D) or peptide (E, F) agrees well with experimental data (dots) and theoretical model (solid lines). **G**. The optimal droplet radius decreases with a faster deactivation rate k_deactivation_. For k_deactivation_ > (2.1 seconds)^-1^, active droplets vanish. **H**. The rate of change in the droplet radius vanishes at the optimal droplet radius. Above, droplets grow by influx of activated material, while below, they shrink by efflux of deactivated material.

This size control mechanism implies that synthetic cells can control the size and number of their organelles by investing chemical fuel in a chemical reaction cycle that activates droplet material in the dilute phase and deactivates it in the droplet phase. We prepared such synthetic cells using an artificial cytoskeleton of an agarose gel that contained precursor, polyanion, and buffer in a solution of perfluorinated oil with DIC. Droplets of similar radii were observed in the synthetic cell, and the total volume of all combined active droplets remained constant, pointing to a steady state (Supplementary Figure 12). We varied the fuel concentration in the perfluorinated oil, which sets the steady-state concentration in the reactors (See Materials and Methods and Supplementary Figure 10D and F). An increase in the steady-state concentration of fuel in the synthetic cell between 2.5 and 10 mM did not change the optimal droplet radii significantly (Figure 3C). Instead, the droplet number density increased (Figure 3B and D). Thus, more fuel produces more droplets of a similar size. In contrast, when we increased the peptide concentration in the solution, the droplet radii increased slightly while the droplet number density remained stable (Figure E and F respectively). Interestingly, we found that for each condition, the radii and their distribution width were independent of the synthetic cell size, even though increasing the size results in larger amounts of droplet material, which was accommodated by creating more droplets (Supplementary Figure 2).

We used this theoretical model to probe these observations and predict the system’s behavior outside of the range of parameters that we could experimentally vary (e.g., the deactivation rate constant or the concentration peptide beyond its solubility). The model confirmed the validity of the observation, *i*.*e*., how the concentrations of fuel and peptide set the optimal droplet radius and the number of droplets in the system (solid lines in Figure 3C-F and Supplementary Figure 13 E-H). We also found that an increasing deactivation rate constant yields smaller droplets until no more droplets are found at a deactivation rate constant corresponding to a half-life of the activated peptide of 2.1 seconds (Figure 3G). Thus, varying the deactivation rate constant is the most effective droplet size control. Biocatalysis can easily achieve this in the context of biomolecular condensates. It would require increased kinase activity in the droplets (deactivation) and increased phosphatase (activation) activity in the cytosol. Such a spatial separation is evidenced in P granules,^30^ stress granules,^31,32^ Cajal bodies.^33^ Our work further adds to the notion that cells can control their organelle size by investing chemical energy in phase separation and spatially separating droplet material activation and deactivation.

Finally, we used the theoretical model to calculate a single droplet’s net influx or efflux as a function of its radius (Figure 3H). These fluxes made it clear that the net efflux increases drastically with increasing radius beyond the optimal radius of 0.75 μm.

## Size control and gravitationally induced fusion drive oscillations

Both experiment and theory demonstrate that synthetic cells can control the size of their active droplets by balancing influx and efflux. This size control mechanism explains the counterintuitive part of the oscillations, *i*.*e*., gravitationally induced fusion leads to droplets at the bottom of the synthetic cell with radii much greater than the optimal radius and thus expel large amounts of deactivated droplet material (Figure 4A). We hypothesize that the expelled, deactivated material creates a concentration gradient from the cell’s bottom to the top. At the bottom, near the large droplets, the deactivated peptide is reactivated and quickly reenters the large droplet. In contrast, at the top of the synthetic cell, the reactivated peptide cannot enter a droplet and thus accumulates. New droplets can only nucleate when sufficient reactivated peptide has accumulated at the top of the synthetic cell. Then, after nucleation, the droplets continue growing while suppressing further nucleation events by locally lowering activated peptide concentration. Once droplets are sufficiently large to outcompete their Brownian motion, they sediment, leading to gravitationally induced fusion at the bottom, restarting the oscillation. The suppression of nucleation is a prerequisite for the oscillations—without it, droplets would continuously be produced at the top, leading to continuous settling of droplets without oscillations.

**Figure 4.**
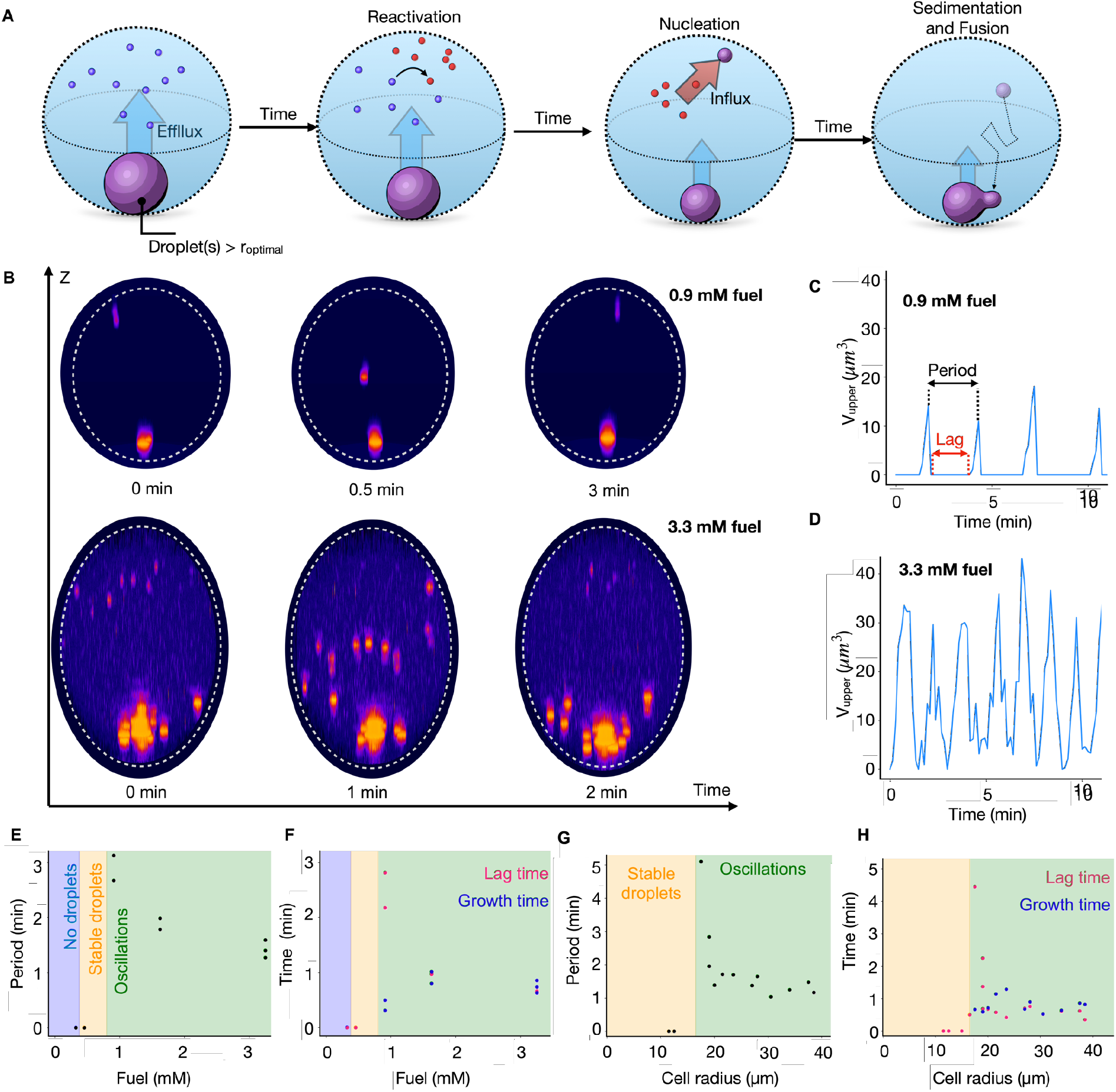
Parameters that control the oscillation. The conditions were 10 mM precursor, 5 mM PSS in 200 mM MES pH 5.3 with 0.5 μM sulforhodamine B as the dye, and 250 mM DIC was added to the oil phase to induce coacervation, except indicated otherwise. **A**. Schematic representation of the events in each oscillation. New droplets nucleate and grow in the upper half of the reactors. Upon growth, these droplets sink down and fuse at the bottom of the reactor while new droplets nucleate in the upper half of the reactor. Grey dotted circles represent the periphery of the reactors, and green dotted circles mark a tracked droplet through the time series. **B**. Snapshots at different times of the side projections of two similarly sized cells when exposed to two different fuel concentrations, 0.9 mM (upper row) and 3.3 mM (lower row). The time evolutions show that the number and the frequency of new droplets waves depends on fuel concentration. Synthetic cells appear elongated due to imaging artifacts. **C**. The combined volume of all the organelles in the upper half at 0.91 mM fuel and **D**. at 3.3 mM fuel. **E**. Period of the oscillations calculated from the dominant frequency determined by FFT as a function of fuel and **G**. cell size. **F**. Lag time between waves of nucleation (pink) and growth time (blue) at different fuel concentrations and **H**. cell radius. In both cases, the lag time follows the same trend as the period, suggesting that the lag strongly influences the frequency of the oscillations.

As the reactivation of peptide and the consequent droplet nucleation of droplets requires fuel, there should be a minimum fuel concentration to trigger the oscillations. Below this concentration, insufficient droplet material is reactivated at the top of the reactor to induce nucleation and, thus, the next wave of droplets. Instead, it will diffuse back to the large droplet at the bottom of the synthetic cell. We found that the minimum fuel concentration needed to induce oscillations was 0.9 mM (Figure 4E). At 0.9 mM, we observed a single nucleation event at the top of the synthetic cell (Figure 4B top, Supplementary Movie 5). This droplet sediments to the bottom and, finally, fuses with the big droplet, starting the cycle again. Below this fuel concentration, we found stable droplets at the bottom that did not oscillate (Figure 4E). If we further decreased the fuel concentration, no droplets were found.

In contrast, when we increased the fuel concentration, the synthetic cells produced more droplet material (Figure 4B bottom, Supplementary Figure 14, and Supplementary Movie 6). Moreover, with each wave, more droplets nucleated simultaneously at the top of the synthetic cell, leading to larger droplets at the bottom after fusion. Finally, we noted the period decreased with increasing fuel (Figure 4D-E). Additionally, we can also control the size of the synthetic cell. Like varying the fuel concentration, we observed a threshold for oscillations— synthetic cells with a radius smaller than 16 μm display stable droplets without oscillations. The period decreased rapidly as we increased the radius to 40 μm (Figure 4G). We found that the lag to nucleate new droplets also decreased, *i*.*e*., when the last droplets were fusing, the next wave had already nucleated (Figure 4H). The time this process took seemed to set the lower limit of the oscillation period. To understand the relation between fuel concentration or reactor size and the period of the oscillations, we determined the lag period between the droplets sinking into the bottom part of the cell and the nucleation of new droplets at the top of the synthetic cell (See, for example, Figure 4C). We found that it was as long as 3 minutes for low fuel levels and rapidly dropped with increasing fuel. For example, with 3.3 mM of fuel, the lag period was less than 40 seconds, meaning that new droplets nucleated before the previous wave completely fused. Similarly, the lag could be several minutes for small synthetic cells or very short for large synthetic cells (Figure 4H). Additionally, we calculated the time in which droplets were present in the upper half of the synthetic cell— the growth time—and that it rapidly stabilized (Figure 4F and H). Thus, close to the threshold for oscillations, the period of the oscillations is predominantly determined by the lag phase between droplet settling and re-nucleation of droplets, which is tuneable by the reactor size and the fuel concentration. In contrast, more far away from the threshold for oscillations, growth and lag time approximately contribute equally to the oscillations period (Figure 4F and H).

## Conclusions

In this study, we successfully demonstrated the spontaneous emergence of spatio-temporal oscillations in the number, size, and position of droplets within synthetic cells. These oscillations are driven by a novel size control mechanism combined with gravitationally induced fusion. Our findings reveal that large sedimented droplets shrink by expelling droplet material, which nucleates new droplets at the top of the synthetic cell, sustaining the oscillations through repeated cycles. This oscillatory behavior can continue for hundreds of periods without fatigue and is tunable within the minute range.

Our synthetic cells, which mimic the rhythmic patterns found in natural biological systems, are fueled by a chemical reaction that continuously activates and deactivates peptide molecules. This activation-deactivation cycle is crucial for maintaining the droplets’ dynamic behavior. The system’s ability to control the size and number of droplets through chemical energy investment and spatial separation of activation and deactivation processes brings us closer to creating more life-like materials. The implications of our findings extend beyond the immediate study of oscillations in synthetic cells. The ability to design and control rhythmic patterns in synthetic cells opens new avenues for advancing synthetic biology and systems chemistry.

## Acknowledgments

The BoekhovenLab is grateful for support from the TUM Innovation Network - RISE funded through the Excellence Strategy. This research was conducted within the Max Planck School Matter to Life, supported by the German Federal Ministry of Education and Research (BMBF) in collaboration with the Max Planck Society. Funded by the Deutsche Forschungsgemeinschaft (DFG, German Research Foundation) – Project-ID 364653263 – TRR 235, under Germany’s Excellence Strategy - EXC-2094 – 390783311 and the European Research Council (ERC starting grant 852187). The Weber group acknowledges funding by the European Research Council (ERC) under the European Union’s Horizon 2020 research and innovation program (Fuelled Life with Grant agreement No 949021).

## Materials and Methods

### Materials

Reagents and buffers were purchased from Sigma-Aldrich and used without further purification unless otherwise indicated. HPLC-grade acetonitrile (ACN) was purchased from VWR. The peptide Ac-F(RG)_3_N-NH_2_ (99%) was purchased from CASLO Aps. 2 w% 008-Fluorosurfactant in Novec 7500 was purchased from RAN Biotechnologies. Novec 7500 was purchased from Iolitec.

### Manual solid-phase peptide synthesis under controlled heating

AcF(RG)_3_D-OH was synthesized on a 0.25 mmol scale using aspartic acid preloaded Wang resin (0.25 mmol, 0.67 mmol g-1). The reaction vessel was connected to a nitrogen line, and the waste flask to a water pump. The resin was swell in DMF for 30 min with medium N_2_ bubbling at room temperature. The peptide synthesis was performed at 68 °C.

Before each coupling step, the *N*-terminal Fmoc-protecting group was cleaved using 5 mL of a 0.2 M HOBt 5% (w/v) solution of piperazine in DMF. The reaction mixture was stirred with an N2 stream for 1 minute and 5 min. The deprotecting solution was removed, and the resin was washed with DMF. For each coupling, 3 eq. of amino acid were used together with 2.8 eq. of HCTU and 6 eq. of DIPEA. HCTU and DIPEA were added to the amino acid and vortexed until dissolved. The coupling solution was added to the resin and stirred with an N2 stream for 7 min. After the coupling, the solution was drained, and the resin was washed with DMF. The deprotection, washing, coupling, and washing cycle was carried out for each amino acid. After the last coupling, the peptide was acetyl end-capped at room temperature. Therefore, a final deprotection was conducted; the resin was washed at room temperature, and 6 eq. of acetic anhydride and 6 eq. of DIPEA in DMF solution were added to the resin and stirred with N2 stream for 10 min at RT. The resin was washed with DMF and DCM. A cleavage solution consisting of 2.5 % MQ-water, 2.5 % TIPS, and 95 % TFA was prepared to cleave the peptide from the resin. It was added to the resin and agitated for two hours at RT. The cleavage solution was collected by filtration, and the resin was washed with DCM. The solvents were removed by co-distillation under reduced pressure using a rotary evaporator (Hei-VAP Core, VWR). The crude peptides were dissolved in H2O:ACN (80:20) and purified using a reversed-phase preparative HPLC (Agilent 1260 Infinity ll setup, Agilent InfinityLab ZORBAX SB-C18 column 250 mm×21.2 mm, 5 μm particle size, linear gradient of ACN from 2% to 98% and water, both with 0.1% TFA, flow rate 20 mL min^-1^, retention time = 6 min) The peptide was lyophilized (Christ Freeze Dryer Alpha 2-4 LDplus, VWR) and stored at -20 °C. It was characterized by electrospray ionization mass spectrometry (ESI-MS, Thermo Scientific, LCQ Fleet ION Trap Mass Spectrometer) in positive mode (Mass calc: 963.05 g/mol; Mass observed: 963.30 [Mw]+), analytical HPLC (Thermofisher Ultimate 3000, 250 mm × 4.8 mm Hypersil Gold C18 column, linear gradient of acetonitrile 2% to 98% and water with 0.1% TFA) and ^1^H NMR. Characterization data was in agreement with already published data.^20^

### Kinetic model

We used a kinetic model to predict the evolution of the product concentration over time and to model the steady-state product concentration at different conditions previously reported by Bergmann et al.^19^ for the same peptide and carbodiimide. Supplementary Method 1 describes the model briefly. The rate constants used in this work are shown in Supplementary Table 1.

### Steady-state kinetics HPLC sample preparation

200 μL of the aqueous phase was prepared by adding the appropriate volumes of the precursor stock solution and diluting them in 0.2 M MES buffer pH = 5.3 to achieve a final concentration of 8-16 mM precursor. To simulate the gel layer system, 200 μL of pure DIC were added on top and the aqueous phase was gently stirred. To mimic the synthetic cell container experiments, 1 mL of oil phase was prepared by adding DIC to perfluorinated oil (Novec 7500) to a final concentration of 50-500 mM. Then, the aqueous phase containing the precursor (200 μL) was added on top and the solution was vigorously stirred to form an emulsion. In both cases, 100 μL of the aqueous solution was pipetted from the solution and mixed with 100 μL of amylamine 200 mM to stop the reaction. Next, we performed an extraction step by adding 200 L of chloroform to remove DIC, which facilitates HPLC analysis. Finally, the aqueous phase was collected and injected without further dilution. All data points correspond to independent samples and were performed in triplicate (N=3).

### Nuclear magnetic resonance spectroscopy (NMR)

^1^H-NMR spectra were recorded with 16 scans at room temperature on a Bruker AVHD 300 spectrometer. Chemical shifts are reported in parts per million (ppm) relative to the signal of the reference compound TMPS (∂ = -0.017 ppm). All measurements were performed in duplicate (N = 2) at 25 °C. Concentrations were determined from the peak integrals, which were compared with the TMPS reference. The chemical shifts of the compared signals were ^1^H NMR (300 MHz, TMPS): TMPS δ (ppm) = 0 (s, 9H, CH_3_), DIC δ (ppm) = 1.26-1.14 (d, 12H, CH_3_), DIU δ (ppm) = 1-1.12 (d, 12H, CH_3_).

### Steady-state kinetics NMR sample preparation

400 μL of the aqueous phase containing 14 mM precursor and 50 mM TMPS in 0.2M MES pH 5.3 were prepared by dissolving the dry powder of TMPS and diluting the precursor stock. To start the reaction, 400 μL of DIC were added on top and the system was let to evolve for a certain amount of time. After that, 300 μL of the aqueous phase were mixed with 300 μL 640 mM borate buffer pH 10 containing 20% D_2_O to quench the reaction by vortexing. All datapoints correspond to independent samples and were performed in duplicate (N=2).

### Determination of DIC K_p_

To determine the partition coefficient between the perfluorinated oil and 0.2M MES buffer, pH 5.3 we dissolved 50 mM TMPS as reference in 500 μL buffer. 2.5 mL of perfluorinated oil containing 0.15-1 M DIC were added on top and samples were stirred vigorously to form an emulsion. After 4h, the two phases were allowed to separate and 300 μL of the aqueous solution were quenched with 300 μL 640 mM borate buffer pH 10 containing 20% D_2_O. DIC and DIU in the solution were determined by ^1^H NMR spectroscopy as described above. DIC concentration in the aqueous phase was plotted against DIC concentration in the oil phase to obtain K_p_. Values are collected in Supporting Figure 10.

### Determination solubility of DIC in 0.2 M MES buffer at pH=5.3

DIC solubility was determined by ^1^H NMR spectroscopy. Samples were prepared by adding 500 μL of DIC on top of 500 μL 0.2 M MES buffer pH 5.3 and stirring gently for 6h. After, 300 μL of the aqueous phase were quenched with 300 μL of 640 mM borate buffer pH 10 20% D_2_O and measured as described above. Values are collected in Supporting Figure 10.

### High-performance liquid chromatography (HPLC)

High-pressure liquid chromatography was performed using analytical HPLC (Thermofisher Ultimate 3000) with a Hypersil Gold C18 column (250 mm × 4.8 mm). Separation was performed using a linear gradient of acetonitrile (2% to 85%) and water with 0.1 vol% TFA, and the chromatogram was analyzed using detector at 220 nm. The data was collected and analyzed with the Chromeleon 7 Chromatography Data System Software (Version 7.2 SR4). All measurements were performed in triplicate (N = 3) at 25 °C. Details about the solvent gradient used to determine peptide and active peptide concentration are shown in Supplementary Table 4.

### Method to sustain steady-state in confinement

We produced surfactant-stabilized water in oil droplets using 1 w% 008-FluoroSurfactant in 3 M HFE7500 as the oil phase. To form containers of varying size, 5 μL of the aqueous solution were added to 50 μL of the oil phase in a 200 μL Eppendorf tube. Snipping of the centrifugal tube resulted in the formation of containers with a random size distribution. The microreactors were imaged at the confocal microscope in untreated observation chambers consisting of a 24 mm × 60 mm glass cover slide and a 16 mm × 16 mm glass cover slide that were separated by two slices of double-sided sticky tape and sealed with two-component glue. For experiments that require hindered fusion, the aqueous solution contained 8-16 mM precursor, 8.6-17 mM PSS and 0.2 μM sulforhodamine B in 0.5 w/v % agarose 200 mM MES at pH 5.3 and the oil phase contained 0-500 mM DIC. Both solutions were kept at 45 °C until they were mixed to ensure water in oil droplet formation before agarose gelation.

In free fusion experiments, the aqueous solution contained 10 mM precursor, 5 mM PSS and 0.5 μM in 200 mM MES at pH 5.3 and the oil phase contained 0-250 mM DIC.

In both cases, samples were imaged after 30 minutes to guarantee that the steady-state concentrations were already reached.

### Method to sustain steady-state in bulk

We prepared a layer of lower melting point (LMP) agarose gel (0.5 w/v % except otherwise stated) containing 14 mM precursor, 15 mM PSS and 0.2 μM sulforhodamine B by heating a 0.75 w/v % LMP agarose 0.2 M MES buffer pH 5.3 stock solution to 45 °C and adding the appropiate volumes of precursor, PSS and sulforhodamine B stock solutions, all in 0.2 M MES buffer pH 5.3. 15 μL of the resulting solution were transferred to an Ibidi μ-Slide 15 Well 3D Glass Bottom previously passivated with PVA while still warm and let to jellify for 10 minutes. This resulted in a gel layer of approximately 1 mm thickness, which ensured no fuel gradients. To start the reaction, we added 15 μL of DIC on top and started imaging immediately.

For prenucleated droplets, the same procedure was followed but 20 mM EDC were added to the aqueous solution while still liquid to trigger coacervate formation which allowed droplet fusion for a few minutes resulting in a wide size initial distribution. Next, the solution was transferred to a well and, after 3 minutes, the DIC layer was added and imaging started.

### Passive droplets preparation into a gel matrix

To study the effect of the gel on droplet size, we performed control experiments –passive droplets– where 1.4 mM of precursor were substituted by a non-hydrolysable mimic of our product (product*, AcF(RG)_3_N-NH_2_). In this case, a solution of 12.6 mM precursor, 1.4 mM product*, 15 mM PSS and 0.2 μM sulforhodamine B in 0.5 w/v % agarose 0.2 M MES pH 5.3 was prepared while the agarose solution was still warm and transferred to an Ibidi μ-Slide 15 Well 3D Glass Bottom previously passivated with PVA. Sample imaging was started after 3 minutes.

### Confocal fluorescence microscopy

A Stellaris 5 confocal microscope with a 63x oil immersion objective (1.4 NA) was used to image the samples. We used sulforhodamine B as droplet dye, excited the fluorophore at 561 nm and imaged at 570-650nm with a HyD detector. The pinhole was set to 1 Airy unit. Typically, pixel size was set to 103 nm and image size was set to 512 px x 512 px. For hindered fusion experiments in bulk, we imaged a 6 μm in thickness layer with a z-step of 0.36 μm and a time step of 30 s. The same z-step was applied to experiments under confined conditions, but z-size was modified to image the whole container. In free fusion experiments, time step was set to 9-11 s and z-step to 4 μm. All experiments were performed in triplicate (N=3).

### Image analysis

All images were processed and analyzed using ImageJ (Fiji).^34^ First, a maximum intensity z-projection was obtained. Then, we applied a Gaussian Blur (sigma = 2 px) followed by thresholding with the Otsu algorithm. Next, we used the analyze particles plugin (size = 0.1 μm^2^-infinity, circularity = 0.8-1) to obtain the area and the xy position of each droplet at each timepoint as well as the droplet number in each frame. To finish, we used a modified version of Simple Tracker,^35^ based on MATLAB^36^, to reconstruct each droplet trajectory using the Hungarian linker with 1 μm as maximum linking distance and a maximum gap of 3 frames. Droplet radius and volume was calculated from the area assuming a spherical droplet. Total volume of droplets was calculated as the sum of each droplet’s volume. The kymograph was generated using ImageJ (Fiji). Therefore, a Z-stack time series was converted into an interpolated 3D projection along the Y-axis. The kymograph was obtained along a vertical line with a width of 400 pixels using the plugin KymographBuilder.

### Determination of the period, lag time, and growth time in oscillating synthetic cells

Wave parameters were determined using the total volume of droplets in the upper half of the cell over time. The signal was denoised using a rolling average (window = 3. The period was calculated from the dominant frequency which was determined by performing a fast Fourier transformation (FFT) using Scipy^37^ in Python 3.4 after normalizing the signal to μ=0 and σ=1. The lag time was measured as the time between the start and the end of a trough. The value was obtained from the average of all cycles for a given signal. Growth time was calculated by subtracting the lag time from the period.

### Statistics

Statistical analyses were performed using Excel. Experimental data were analyzed using a two-tailed t-test for two samples assuming unequal variances. P < 0.05 was considered statistically significant. The exact sample size n, alpha level α, and P-value for each test are given in the source data or Supplementary Information.

## Notes

### Competing Interest Statement

The authors have declared no competing interest.

