## Supplementary Information for "Size control and oscillations of active droplets in synthetic cells"

**References** **25**

### Supplementary Methods

#### Supplementary Method 1 : Kinetic model

The reaction cycle in a homogeneous system is described in a kinetic model according to the following mechanism:

$\mathrm{F} \xrightarrow{k_0} \mathrm{W}$

$\mathrm{Ac} + \mathrm{F} \xrightarrow{k_1} \mathrm{I}$

$\mathrm{I} \xrightarrow{k_{An}} \mathrm{An} + \mathrm{W}$

$\mathrm{I} \xrightarrow{k_{Ac}} \mathrm{Ac} + \mathrm{W}$

$\mathrm{An} \xrightarrow{k_2} \mathrm{Ac}$

Ac is the dicarboxylic acid, F is the fuel, I is the intermediate O-acylurea, W is the waste, and An is the anhydride. The mechanism translates into the following set of differential equations:

$$
\begin{aligned}
\frac{d[Ac]}{dt} &= -k_1 \cdot [Ac] \cdot [F] + k_2 \cdot [An] + k_{Ac} \cdot [I] \\
\frac{d[F]}{dt} &= -k_1 \cdot [Ac] \cdot [F] - k_0 \cdot [F] \\
\frac{d[W]}{dt} &= k_0 \cdot [F] + k_{Ac} \cdot [I] + k_{An} \cdot [I] \\
\frac{d[An]}{dt} &= k_{An} \cdot [I] - k_2 \cdot [An] \\
\frac{d[I]}{dt} &= k_1 \cdot [Ac] \cdot [F] - k_{An} \cdot [I] - k_{Ac} \cdot [I]
\end{aligned} \tag{1}
$$

We then applied steady-state approximation to obtain:

$$
\frac{d[I]}{dt} = k_1 \cdot [Ac] \cdot [F] - k_{An} \cdot [I] - k_{Ac} \cdot [I] \approx 0 \implies [I] \approx \frac{k_1 \cdot [Ac][F]}{(K+1)} \tag{2}
$$

We called $k_{Ac}/k_{An} = K$ and used the relation above in the set of differential equations to obtain:

$$
\begin{aligned}
\frac{d[Ac]}{dt} &= -k_1 \cdot [Ac] \cdot [F] + k_2 \cdot [An] + \frac{K \cdot k_1 \cdot [Ac] \cdot [F]}{(K+1)} \\
\frac{d[F]}{dt} &= -k_1 \cdot [Ac] \cdot [F] - k_0 \cdot [F] \\
\frac{d[W]}{dt} &= k_0 \cdot [F] + k_1 \cdot [Ac] \cdot [F] \\
\frac{d[An]}{dt} &= \frac{k_1 \cdot [Ac] \cdot [F]}{(K+1)} - k_2 \cdot [An]
\end{aligned} \tag{3}
$$

Experimental data were fit to the equation system (3) using a custom program in Python 3 (kinmodel, https://github.com/scotthartley/kinmodel) previously published by the group of Hartley and applied to similar systems [4]. To calculate concentrations of precursor and product under continuous fueling, the change in fuel concentration was set to 0 ($d[F]/dt = 0$).

Next we give the details of the theoretical model used to describe the experimental trends of the average radius with system size and fuel concentration. As the experimental setting (chemical compounds, reaction cycle, etc.) is similar to Ref. [1], we also use the theoretical model described in the Supplementary Information of this reference. For clarity, we review the key equations and methods used to obtain the results given in this paper. We also give details on the numerical method used to integrate the equations of our model, and provide estimates on the inter-droplet distance, the reaction-diffusion length scales, and the sedimentation speed.

#### Supplementary Method 2 : Thermodynamic model for the experimental phase diagram

In our theoretical model, we consider an effective ternary mixture where the effects of fuel, waste, and polyanion are accounted for in an implicit manner. The effective components in the model are the solvent, the precursor $A$, and the product $B$. For these components and a few experimental conditions, the equilibrium concentrations in the dense and the dilute phases were determined experimentally (see Ref. [1]). These experimental points gave a rough idea of the underlying binodal line that separates the domain of phase separation and a homogeneous mixture in the phase diagram. To continuously interpolate these data points and obtain insights into the molecular interactions among the components, we describe the experimental data points using a theoretical model for the binodal line in the phase diagram.

To calculate the theoretical phase diagram, we consider a mean-field model with the following free energy density:

$$f(c_A, c_B) = k_B T \left[ \sum_{i=A,B,S} c_i \log(\nu_i c_i) + \sum_{ij=AB,AS,BS} \chi_{ij} (\nu_i c_i)(v_j c_j) \right] , \tag{4}$$

where $c_A$, $c_B$ and $c_S$ denote the number concentrations of components $A$, $B$, and solvent, respectively. The molecular volumes of components $A$, $B$ and the solvent are $\nu_A$, $\nu_B$ and $\nu_S$, and their products with the respective number concentrations, $(\nu_i c_i)$, are volume fractions. We consider an incompressible mixture where all molecular volumes are constants. In this case, we can describe the thermodynamics of the mixture by three volume fractions or, equivalently by the three concentrations $c_A$, $c_B$ and $c_S$. These concentrations obey $c_S = 1/\nu_S - r_A c_A - r_B c_B$ with $r_i = \nu_i/\nu_S$ as the molecular volume ratios. Using this relationship, we can substitute the solvent concentration and recast the free energy density $f(c_A, c_B)$ as a function of $c_A$ and $c_B$. This free energy also depends on the parameters $r_A$ and $r_B$, as well as the parameters describing the inter-molecular interaction: $\chi_{AB}$, $\chi_{AS}$ and $\chi_{BS}$. All such parameters were determined by the comparison between the experimentally measured equilibrium concentrations and the theoretical thermodynamic model based on the free energy density (Eq. (4)).

At phase equilibrium, the exchange chemical potentials $\mu_i = \partial f / \partial c_i$ $(i = A, B)$ of components $A$ and $B$ and the osmotic pressure $\Pi = -f + \sum_{i=A,B} c_i \mu_i$ are balanced between the phases I and II:

$$\mu_A\left(c_A^{\mathrm{I}}, c_B^{\mathrm{I}}\right) = \mu_A\left(c_A^{\mathrm{II}}, c_B^{\mathrm{II}}\right) , \tag{5a}$$

$$\mu_B\left(c_A^{\mathrm{I}}, c_B^{\mathrm{I}}\right) = \mu_B\left(c_A^{\mathrm{II}}, c_B^{\mathrm{II}}\right) , \tag{5b}$$

$$\Pi\left(c_A^{\mathrm{I}}, c_B^{\mathrm{I}}\right) = \Pi\left(c_A^{\mathrm{II}}, c_B^{\mathrm{II}}\right) . \tag{5c}$$

The set of points $(c_A, c_B)$ satisfying these three conditions make up the binodal line in the phase diagram. In this work, we find a good agreement with the experimental data when using the following parameters: $r_A = 30$, $r_B = 60$, $\chi_{AB} = -0.18$, $\chi_{AS} = 0.77$, $\chi_{BS} = 0.81$. These parameters suggest that the interactions between the precursor $A$ and product $B$ are effectively attractive, while each component dislikes the solvent, giving rise to phase separation from the solvent, also in the absence of the respective other component.

#### Supplementary Method 3 : Continuous theory for the dynamics of concentration fields

To capture the dynamics of many droplets in an active emulsion, we describe the spatio-temporal evolution of the concentration fields $c_A(\boldsymbol{x}, t)$ and $c_B(\boldsymbol{x}, t)$, where $\boldsymbol{x}$ is position and $t$ time. To this end, we consider the total free energy:

$$F = \int_V d^3r \left[ f(c_A, c_B) + \frac{\kappa_A}{2}(\nabla c_A)^2 + \frac{\kappa_B}{2}(\nabla c_B)^2 \right] , \tag{6}$$

where $V$ is the volume of the system and $f(c_A, c_B)$ is mean-field free energy density given in Eq. (4). The coefficients $\kappa_A$ and $\kappa_B$ characterize the free energy costs for spatial gradients of the components $A$ and $B$. They determine the width of the droplet interface and are related to the interfacial tensions.

The diffusion of different components in a non-dilute mixture is driven by spatial gradients in the respective chemical potentials, while the chemical reactions minimize the difference in chemical potentials between reactants and products. In our reduced model, the reacting components are $A$ and $B$ that convert into each other following an effective uni-molecular scheme:

$$\mathrm{A} \overset{\mathrm{c_F}}{\rightleftharpoons} \mathrm{B} \, . \tag{7}$$

This uni-molecular reaction has the following conserved quantity:

$$c = V^{-1} \int_V d^3x \, \left(c_A\left(\boldsymbol{x}, t\right) + c_B\left(\boldsymbol{x}, t\right)\right) \,, \tag{8}$$

where $V$ is the system volume.

The presence of fuel of concentration $c_F$ (Eq. (7)) strongly alters the balance between the reacting components $A$ and $B$, i.e., from almost all reacting components in state $A$ at thermodynamic equilibrium to similar amounts of both $A$ and $B$ for most studied experimental conditions. Thus, the thermodynamic pathway from $A$ to $B$ is negligible compared to the pathway mediated by the hydrolysis of fuel components to waste. The maintenance of fuel concentrations the experiments suggests neglecting the thermodynamic pathway $A \to B$ and describing the fuel-driven activation pathway with a rate coefficient $k_{\mathsf{a}}$ that requires a non-zero fuel concentration $c_F$:

$$k_{A\to B} = k_{\mathsf{a}} \, c_F \,. \tag{9}$$

Here, the subscript 'a' refers to the fuel-driven chemical pathway of activation. The deactivation pathway occurs spontaneously. Since the fraction $k_{\mathsf{a}} \, c_F / k_{B\to A}$ is not governed by the free energy of the components $A$, $B$, and solvent, detailed balance of chemical reaction rates is broken. In other words, the system is maintained away from thermodynamic equilibrium due to the maintained fuel in the system. As the system can also phase separate forming multiple droplets, such a system is referred to as chemically active emulsion [3]. For simplicity, the rate coefficients are considered to be constants that we estimated from experimental measurements (details see Ref. [1]). The kinetic equations for the fields $c_A(\boldsymbol{x}, t)$ and $c_B(\boldsymbol{x}, t)$ then read ($i = A, B$):

$$\partial_t c_i = \nabla \cdot \left(\Lambda_i \nabla \mu_i\right) + k_{j\to i} c_j - k_{i\to j} c_i \,, \tag{10a}$$

where $k_{j\to i}$ is the rate coefficient from component $j$ to $i$. The chemical potentials can be calculated via

$$\mu_i = \delta F / \delta c_i \tag{10b}$$

with the functional derivative $\delta F / \delta c_i$ of the total free energy given in Eq. (6). Moreover, $\Lambda_A$ and $\Lambda_B$ are mobilities coefficients of components $A$ and $B$, respectively. They depends of concentrations, $\Lambda_A = \Lambda_A^0 c_A (1 - \nu_S c_S)$ and $\Lambda_B = \Lambda_B^0 c_B (1 - \nu_S c_S)$, where $\Lambda_A^0$ and $\Lambda_B^0$ are constants. The mobilities are related to the diffusion coefficients in each of the phases through

$$D_i^\alpha = \Lambda_i(c_i^\alpha) f''(c_i^\alpha) \,, \tag{10c}$$

with $\alpha =$ I,II. The diffusion coefficients were estimated experimentally for both components in both phases, which sets the values of the constants $\Lambda_A^0$ and $\Lambda_B^0$.

##### Numerical solutions to the continuous dynamic theory

To solve dynamic equations (10) numerically, we used a discrete spatial grid with a grid resolution of around 0.067 $\mu m$. The grid resolution was chosen such that the droplet interface was represented by at least five grid points. This was achieved by an appropriate choice of the free energy costs $\kappa_i$.

For time integration we used a semi-implicit Fourier spectral method [5]. In this method, we solved for the concentration fields in Fourier space. The Fourier transform of the coupled partial differential equations given in (10) gives coupled ordinary differential equations, which were solved using the semi-implicit scheme discussed in [5]. Taking the inverse Fourier transform, we obtain the spatial concentration profiles $c_i(\boldsymbol{x}, t)$. We used the fast Fourier transform package from Python, which naturally implements a periodic boundary condition for our system. The parameters used for the spatial simulations are given in Table 3.

We used the spatial profiles $c_i(\boldsymbol{x}, t)$ to calculate each droplets' radius $R_j$ and the number of droplets in the system volume $V$. A droplet is bound by its interface which we defined as the line at which the equal concentration contours takes the value $c = (c_{\text{in}}^{\text{eq}} + c_{\text{out}}^{\text{eq}})/2$, where $c$ is the total concentration of $A$ and $B$. For large systems containing many droplets we determined the droplet size distribution from which the average radius and the standard deviation were calculated.

The figure (11) shows the kinetics of average radius and standard deviation of the droplet size distribution obtained using the continuous theory.

##### Mapping the conserved variable from experiments to simulations

Since we run the simulations in two spatial dimensions, the mapping between experimentally measured concentrations and the conserved variable in the simulations is slightly non-trivial. We know that the concentration ($c$) and volume fraction ($\phi$) in three dimensions are related as $c = \phi/\nu$, where $\nu$ is the molecular volume. We map the volume fraction $\phi$ to the area fraction $\phi_{2D}$ as follows :

We use a mapping which preserves the ratio of the droplet radius to the inter droplet distance in three as well as two dimensions. We denote the half of the inter-droplet distance by $R_{dd}$ and the droplet radius by $R$. We call each droplet and the surrounding region within a distance $R_{dd}$ as a subsystem. In three dimensions, if the average volume fraction is $\phi$, then the droplet volume and the subsystem volume are related by

$$\phi = \frac{(4\pi/3)R^3\phi^{\text{I}} + (4\pi/3)(R_{dd}^3 - R^3)\phi^{\text{II}}}{(4\pi/3)R_{dd}^3} \ ,$$

as per the conservation of material, or

$$\left(\frac{R}{R_{dd}}\right)^3 = \frac{\phi - \phi^{\text{II}}}{\phi^{\text{I}} - \phi^{\text{II}}} \ , \tag{11}$$

as dictated by conservation of material. Here we assume that the gradients in volume fraction are weak so we can use the equilibrium volume fractions $\phi^{\text{I}}$ and $\phi^{\text{II}}$ to represent the volume fractions inside and outside the droplet. Similarly, in 2D we can write

$$\left(\frac{R}{R_{dd}}\right)^2 = \frac{\phi_{2D} - \phi^{\text{II}}}{\phi^{\text{I}} - \phi^{\text{II}}} \ , \tag{12}$$

where $\phi_{2D}$ is the average area fraction. Now we demand that $R/R_{dd}$ be equal in two and three dimensions, which gives us the relation

$$\phi_{2D} = (\phi_{3D} - \phi^{\text{II}})^{\frac{2}{3}}(\phi^{\text{I}} - \phi^{\text{II}})^{\frac{1}{3}} + \phi^{\text{II}} \ . \tag{13}$$

This gives a relation between total conserved material concentration and the conserved variable in our simulations.

#### Supplementary Method 4 : Sharp interface model for the steady state of active droplets

Since the width of the interface is much smaller than the typical droplet size considered in the experiments, we considered the limit of a sharp interface. In this limit, we study the variation in the steady state radius as a function of the fuel concentration $c_F$ and the conserved quantity $c_{\text{tot}}$ of the effective chemical reaction (7); see Eq. (8) for the definition of the conserved quantity. Moreover, note that the fuel enters implicitly via Eq. (9).

In this limit of a sharp interface, we can calculate the growth rate of droplets $dR/dt$ as a function of the radius $R$. To this end, we consider a spherical droplet in a spherically symmetric container of volume $V_{\text{sys}} = (4\pi/3)R_{\text{sys}}^3$, where $R_{\text{sys}}$ is the radius of the container. At the interface, we impose local phase equilibrium (Eqs. (5)). Away from the interface, inside and outside the droplets, we expand up Eqs. (10) to the first order in the concentration fields in each phase around the respective phase equilibrium concentrations at the interface. The results are linear reaction-diffusion equations, where diffusion is driven by concentration gradients and the chemical reactions follow a first-order reaction kinetics:

$$\partial_t c_i^\alpha = D_i^\alpha \nabla^2 c_i^\alpha + k_{i\to j}^\alpha c_j^\alpha - k_{j\to i}^\alpha c_i^\alpha \ , \tag{14}$$

where $\alpha = \text{I}, \text{II}$, where I denotes the droplet phase and II indicates the phase outside the droplet. Note that since diffusion coefficients $D_i^\alpha$ and linear reaction rate coefficients $k_{i\to j}^\alpha$ are evaluated at the expansion points of the equilibrium concentrations at the interface, i.e., they depend on the respective phases I and II.

At steady state ($\partial_t c_i^\alpha = 0$), the concentration profiles can be determined as a function of the radial coordinate $r$ from Eq. (14). To calculate the stationary radius, we impose no flux boundary conditions at the droplet centre at $r = 0$ and at the system boundary $r = R_{\mathsf{sys}}$. Note that for calculating the growth rate $\dot{R}$ different boundary conditions have to be considered; see below. At the droplet interface at $r = R$, we impose $c_A^\alpha(R) = a^\alpha$, $c_B^\alpha(R) = b^\alpha$ with $\alpha =$ I,II. For these boundary conditions, the stationary solutions inside the droplet (phase I) read

$$c_A^{\mathsf{I}}(r) = \frac{a^{\mathsf{I}}\rho_k^{\mathsf{I}} - b^{\mathsf{I}}}{\rho_D^{\mathsf{I}} + \rho_k^{\mathsf{I}}}\frac{R\sinh(r/\lambda^{\mathsf{I}})}{r\sinh(R/\lambda^{\mathsf{I}})} + \frac{a^{\mathsf{I}}\rho_D^{\mathsf{I}} + b^{\mathsf{I}}}{\rho_D^{\mathsf{I}} + \rho_k^{\mathsf{I}}}\,, \tag{15a}$$

$$c_B^{\mathsf{I}}(r) = \rho_D^{\mathsf{I}}\frac{b^{\mathsf{I}} - a^{\mathsf{I}}\rho_k^{\mathsf{I}}}{\rho_D^{\mathsf{I}} + \rho_k^{\mathsf{I}}}\frac{R\sinh(r/\lambda^{\mathsf{I}})}{r\sinh(R/\lambda^{\mathsf{I}})} + \rho_k^{\mathsf{I}}\frac{a^{\mathsf{I}}\rho_D^{\mathsf{I}} + b^{\mathsf{I}}}{\rho_D^{\mathsf{I}} + \rho_k^{\mathsf{I}}}\,, \tag{15b}$$

whereas outside the droplet (phase II), the stationary profiles are given by

$$c_A^{\mathsf{II}}(r) = \frac{a^{\mathsf{II}}k_{BA}^{\mathsf{II}} - b^{\mathsf{II}}k_{AB}^{\mathsf{II}}}{k_{BA}^{\mathsf{II}} + k_{AB}^{\mathsf{II}}\rho_D^{\mathsf{II}}}\frac{R}{r}\frac{\sinh(r/\lambda^{\mathsf{II}}) + \Omega\cosh(r/\lambda^{\mathsf{II}})}{\sinh(R/\lambda^{\mathsf{II}}) + \Omega\cosh(R/\lambda^{\mathsf{II}})} - \frac{a^{\mathsf{II}}k_{BA}^{\mathsf{II}} + b^{\mathsf{II}}k_{AB}^{\mathsf{II}}\rho_D^{\mathsf{II}}}{k_{BA}^{\mathsf{II}} + k_{AB}^{\mathsf{II}}\rho_D^{\mathsf{II}}}\,, \tag{15c}$$

$$c_B^{\mathsf{II}}(r) = \rho_D^{\mathsf{II}}\frac{b^{\mathsf{II}}k_{AB}^{\mathsf{II}} - a^{\mathsf{II}}k_{BA}^{\mathsf{II}}}{k_{BA}^{\mathsf{II}} + k_{AB}^{\mathsf{II}}\rho_D^{\mathsf{II}}}\frac{R}{r}\frac{\sinh(r/\lambda^{\mathsf{II}}) + \Omega\cosh(r/\lambda^{\mathsf{II}})}{\sinh(R/\lambda^{\mathsf{II}}) + \Omega\cosh(R/\lambda^{\mathsf{II}})} - \rho_k^{\mathsf{II}}\frac{a^{\mathsf{II}}k_{BA}^{\mathsf{II}} + b^{\mathsf{II}}k_{AB}^{\mathsf{II}}\rho_D^{\mathsf{II}}}{k_{BA}^{\mathsf{II}} + k_{AB}^{\mathsf{II}}\rho_D^{\mathsf{II}}}\,. \tag{15d}$$

We have used reaction-diffusion length scales $\lambda^\alpha = \sqrt{D_A^\alpha D_B^\alpha/(D_A^\alpha k_{AB}^\alpha + D_B^\alpha k_{BA}^\alpha)}$, and $\rho_D^\alpha = D_A^\alpha/D_B^\alpha$ , and $\rho_k^\alpha = k_{BA}^\alpha/k_{AB}^\alpha$, whereas $\alpha = \mathsf{I}, \mathsf{II}$ indicate the phases. The coefficient $\Omega = -(\lambda^{\mathsf{II}}R_{\mathsf{sys}}\cosh[\lambda^{\mathsf{II}}R_{\mathsf{sys}}] - \sinh[\lambda^{\mathsf{II}}R_{\mathsf{sys}}])/(\lambda^{\mathsf{II}}R_{\mathsf{sys}}\sinh[\lambda^{\mathsf{II}}R_{\mathsf{sys}}] - \cosh[\lambda^{\mathsf{II}}R_{\mathsf{sys}}])$ ensures no flux boundary condition at the system boundary at $R_{\mathsf{sys}}$.

The four interface concentrations $a^{\mathsf{I}}$, $a^{\mathsf{II}}$, $b^{\mathsf{I}}$ and $b^{\mathsf{II}}$, and the position of the interface $R$ can be determined using the constraint of local phase equilibrium at the interface. The first three constraints correspond to local phase equilibrium at the droplet interface :

$$\mu_A(a^{\mathsf{I}}, b^{\mathsf{I}}) = \mu_A(a^{\mathsf{II}}, b^{\mathsf{II}})\,, \tag{16a}$$

$$\mu_B(a^{\mathsf{I}}, b^{\mathsf{I}}) = \mu_B(a^{\mathsf{II}}, b^{\mathsf{II}})\,, \tag{16b}$$

$$\Pi(a^{\mathsf{I}}, b^{\mathsf{I}}) = \Pi(a^{\mathsf{II}}, b^{\mathsf{II}}) - \frac{2\gamma}{R}\,. \tag{16c}$$

The conservation of total number of $A$ and $B$ molecules in the system (Eq. (8)) dictates

$$V_{\text{sys}}\, c = \frac{4}{3}\pi \left[ (a^{\text{I}} + b^{\text{I}})R^3 + (a^{\text{II}} + b^{\text{II}})(R_{\text{sys}}^3 - R^3) \right] . \tag{17}$$

At steady state, the interface remains stationary ($dR/dt = 0$). The local conservation of material then implies for the radial component of diffusive fluxes, $j_i^\alpha = \boldsymbol{j}_i^\alpha \cdot \boldsymbol{e}_r$, at the interface:

$$j_i^{\text{I}}(R) = j_i^{\text{II}}(R) , \tag{18}$$

where $\boldsymbol{e}_r$ is the radial unit vector and $j_i^\alpha = -D_i^\alpha \partial_r c_i^\alpha$. The kinetics of the total concentration, $c = c_A + c_B$, is given by

$$\partial_t c^\alpha = -\partial_r (j_A^\alpha + j_B^\alpha) . \tag{19}$$

The chemical reactions do not enter the kinetics of $c^\alpha$ since it is conserved. For the considered, radial symmetric problem, the stationary solution ($\partial_t c^\alpha = 0$) requires that $j_A^\alpha(r) + j_B^\alpha(r) = 0$. The only independent constraint is then Eq. (18) for either $A$ or $B$, since the condition is automatically fulfilled for the other component. The total number of independent constraints is equal to the total number of unknowns. In particular, there are five unknowns, $a^{\text{I}}$, $a^{\text{II}}$, $b^{\text{I}}$, $b^{\text{II}}$ and $R$, and five Eqs. (16a), (16b), (16c), (17) and (18). For any given set of parameters, such five equations are solved numerically, allowing us to determine the stationary radius $R_{\text{stat}}$ as a function of fuel concentration $c_F$ and the conserved material with concentration $c$. The results are given in the main text, Fig.3.

#### Supplementary Method 5 : Average droplet radius as a combined result of thermodynamics and chemical activity

The average droplet radius $\langle R \rangle$ in the emulsion is expressed in terms of the total volume of the dense phase $V_{\text{dense}}$ as

$$\frac{4\pi}{3} \langle R \rangle^3 = V_{\text{dense}}/N , \tag{20}$$

where $N$ is the total number of droplets in the system. For a total volume of the system $V_{\text{tot}}$ that contains $N$ droplets, we can define a droplet number density

$$\rho = N/V_{\text{tot}} . \tag{21}$$

We can now establish a link to the single droplet discussed in the last section. At least for volume considerations, the emulsion with $N$ droplets can be thought as $N$ single droplet systems, where $V_{\text{tot}} = N\, V_{\text{sys}}$. Moreover, we can relate the average radius and the droplet number density as

$$\frac{4\pi}{3} \langle R \rangle^3 \rho = \frac{V_{\text{dense}}}{V_{\text{tot}}} \,. \tag{22}$$

At phase equilibrium, the total volume of the dense phase $V_{\text{dense}}$ is a thermodynamic quantity that is governed by the phase diagram of the system, which is depicted in Fig. 13. Away from phase equilibrium, there is an (often weak) active contribution to the value of $V_{\text{dense}}$ due to the fuel-driven chemical reactions.

For a system relaxing toward thermodynamic equilibrium, an emulsion ripens leading to a decrease of the the droplet number density $\rho$ toward zero in an infinitely big system and toward $1/V_{\text{tot}}$ in a finite system of total volume $V_{\text{tot}}$ (corresponding to a single droplet in the system). Due to the presence of fuel, the system can give rise to a suppression or an arrest of the ripening kinetics giving rise to a non-equilibrium steady state composed of multiple droplets. The corresponding droplet number density should then be determined by thermodynamic as well as kinetic parameters.

We propose that the droplet number density $\rho$ scales with a single length scale $\lambda$ and can be written as

$$\rho = \frac{a}{\lambda^3} \,. \tag{23}$$

Moreover, we assume that this length scale $\lambda = \lambda^{\text{II}}$ is set by the reaction diffusion $\lambda^{\text{II}} = \sqrt{D_A^{\text{II}} D_B^{\text{II}} / (D_A^{\text{II}} k_{AB}^{\text{II}} + D_B^{\text{II}} k_{BA}^{\text{II}})}$ that dictates the concentration profiles outside the droplets. The parameter $a$ is considered to be determined by thermodynamic parameters (interaction parameters, molecular volumes, etc.) and thus it is constant as a function of fuel concentration or total system volume $V_{\text{tot}}$. The droplet number density can be written as:

$$\rho = \tilde{a} \left[ \frac{D_A^{\text{II}} D_B^{\text{II}}}{D_A^{\text{II}} k_{AB}^{\text{II}} + D_B^{\text{II}} k_{BA}^{\text{II}}} \right]^{-3/2} \,. \tag{24}$$

Using the rate estimates obtained in previous studies [1], we write the activation rate as $k_{BA} = 1.7 \times 10^{-4} c_F$ $s^{-1}$ for the fuel concentration expressed in units of mM. Using this relation and substituting the values of the diffusivities and deactivation rate from the Table 2, we can write the droplet number density as a function of $c_F$ as

$$\rho = a \, (120 + 1.7 c_F)^{3/2}, \tag{25}$$

where $a = 10^{-6} \tilde{a} / (D_A^{\text{II}})^{3/2} \; \mu m^{-3} s^{3/2}$ .

Eq. (22) describes how the total volume of the dense phase $V_{\text{dens}}$ and the droplet number density $\rho$

together determine the average radius of the active droplets $\langle R \rangle$. Fig. 13 compares the variation of droplet radius as a function of fuel concentration $c_F$ for two cases: one case where the average radius is obtained as an output of the sharp interface droplet model and the another case where the average radius is determined indirectly using the predicted variation of total condensed volume and droplet number density with fuel. We see agreement between the above two cases. A minor difference arises because them estimate from the total condensed volume does not account for gradients in the concentrations of the two reacting components.

### Supplementary Tables

| Reaction constant | Value |
|---|---|
| $k_0$ | $2.9 \cdot 10^{-4} s^{-1}$ |
| $k_1$ | $1.7 \pm 0.08 \cdot 10^{-1} M^{-1} s^{-1}$ |
| $K$ | $1.34 \pm 1.0$ |
| $k_2$ | $1.2 \pm 0.9 \cdot 10^{-2} s^{-1}$ |

**Supplementary Table 1:** Kinetic parameters determined by the kinetic model as previously reported in [1]

| Quantity | Symbol | Value |
|---|---|---|
| Interaction parameter precursor-product | $\chi_{AB}$ | -0.18 |
| Interaction parameter precursor-solvent | $\chi_{AS}$ | 0.77 |
| Interaction parameter product-solvent | $\chi_{BS}$ | 0.81 |
| Precursor relative molecular volume | $r_A$ | 30 |
| Product relative molecular volume | $r_B$ | 60 |
| Surface tension | $\gamma$ | 300 $\mu N m^{-1}$ |
| Activation rate outside the drop | $k^{\text{II}}_{BA}$ | $0.17 c_F M^{-1} s^{-1}$ |
| Activation rate inside the drop | $k^{\text{I}}_{BA}$ | $0.51 c_F M^{-1} s^{-1}$ |
| Deactivation rate outside the drop | $k^{\text{II}}_{AB}$ | $0.012 s^{-1}$ |
| Deactivation rate inside the drop | $k^{\text{I}}_{AB}$ | $0.012 s^{-1}$ |
| Diffusion coefficient of A outside the drop | $D^{\text{II}}_A$ | $300 \mu m^2 s^{-1}$ |
| Diffusion coefficient of B outside the drop | $D^{\text{II}}_B$ | $300 \mu m^2 s^{-1}$ |
| Diffusion coefficient of A inside the drop | $D^{\text{I}}_A$ | $0.04 \mu m^2 s^{-1}$ |
| Diffusion coefficient of B inside the drop | $D^{\text{I}}_B$ | $0.0073 \mu m^2 s^{-1}$ |

**Supplementary Table 2 :** Table with input parameters used in numerical calculations of the sharp interface model.

The parameter values used in the table above are motivated by the values used in Ref. [1]. The basic ingredients of the system which are the precursor, the product and the fuel are chemically the same as those considered in Ref. [1]. However, the polyanion used in Ref. [1] was a pSS with a molecular weight of $75kDa$, whereas in this work, we used a smaller pSS of molecular weight $17kDa$. Due to this, slightly different values needed to be chosen for the molecular volume ratio $r_B$ and the interaction parameter $\chi_{BS}$. We used the same experimentally measured concentration values as provided in Ref. [1] to determine the parameters $r_A$, $r_B$, $\chi_{AB}$, $\chi_{AS}$, $\chi_{BS}$. The fitted phase diagram is given in Fig. 13.

| Quantity | Symbol | Value |
|---|---|---|
| Interaction parameter precursor-product | $\chi_{AB}$ | -0.18 |
| Interaction parameter precursor-solvent | $\chi_{AS}$ | 0.77 |
| Interaction parameter product-solvent | $\chi_{BS}$ | 0.81 |
| Precursor relative molecular volume | $r_A$ | 30 |
| Product relative molecular volume | $r_B$ | 60 |
| Interface cost for precursor | $\kappa_A$ | 0.4 |
| Interface cost for product | $\kappa_B$ | 0.4 |
| Activation rate outside the drop | $k^{\text{II}}_{BA}$ | $0.34 c_F M^{-1} s^{-1}$ |
| Activation rate inside the drop | $k^{\text{I}}_{BA}$ | $1.02 c_F M^{-1} s^{-1}$ |
| Deactivation rate outside the drop | $k^{\text{II}}_{AB}$ | $0.012 s^{-1}$ |
| Deactivation rate inside the drop | $k^{\text{I}}_{AB}$ | $0.012 s^{-1}$ |
| Mobility coefficient of A | $\Lambda^0_A$ | $300 \mu m^2 s^{-1} k_B T^{-1}$ |
| Mobility coefficient of B | $\Lambda^0_B$ | $300 \mu m^2 s^{-1} k_B T^{-1}$ |

**Supplementary Table 3:** Table with input parameters used in numerical calculations of the continuous dynamic theory.

| Time (min) | Flow (mL/min) | $H_2O+$ TFA $0.1\%$ | ACN + TFA $0.1\%$ |
|---|---|---|---|
| 0 | 1.00 | 98% | 2% |
| 17 | 1.00 | 15% | 85% |
| 22 | 1.00 | 15% | 85% |
| 22.5 | 1.00 | 0% | 100% |
| 27 | 1.00 | 0% | 100% |
| 28 | 1.00 | 98% | 2% |
| 30 | 1.00 | 98% | 2% |

**Supplementary Table 4 :** HPLC Linear gradient solvent for peptide precursor and product determination.

### Supporting Figures

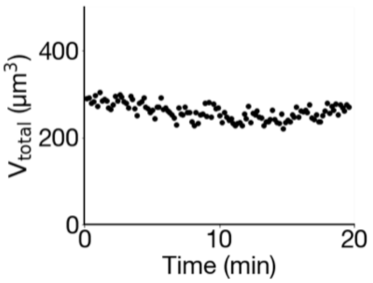

**Supporting Figure 1. Total volume of droplets over time in a synthetic cell showing oscillations.** Conditions were 10 mM peptide, 5 mM PSS expressed in negative charges, 0.5 $\mu$M sulforhodamine B, 200 mM MES at pH 5.3 with 250 mM fuel in the oil phase as fuel. As a result of the steady-state, the volume of droplets remains constant over time

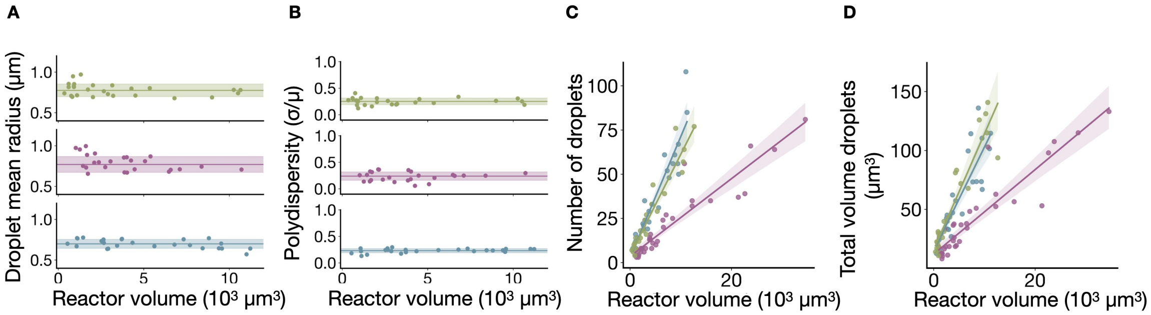

**Supporting Figure 2. Synthetic cell size influence on droplet properties and quantity.** Droplet mean radius (**A**) and polydispersity (**B**) are independent of reactor volume. Number of droplets (**C**) and total volume of droplets (**D**) increase linearly with reactor volume and depend on the concentration of product achieved (in blue, 8mM precursor + 500 mM DIC, in purple 4 mM precursor + 100 mM DIC and in green, 14 mM precursor + 500 mM DIC). Points represent the value obtained for a single synthetic cell of a given size from three independent samples, solid lines the linear fitting and shadowing to the CI.

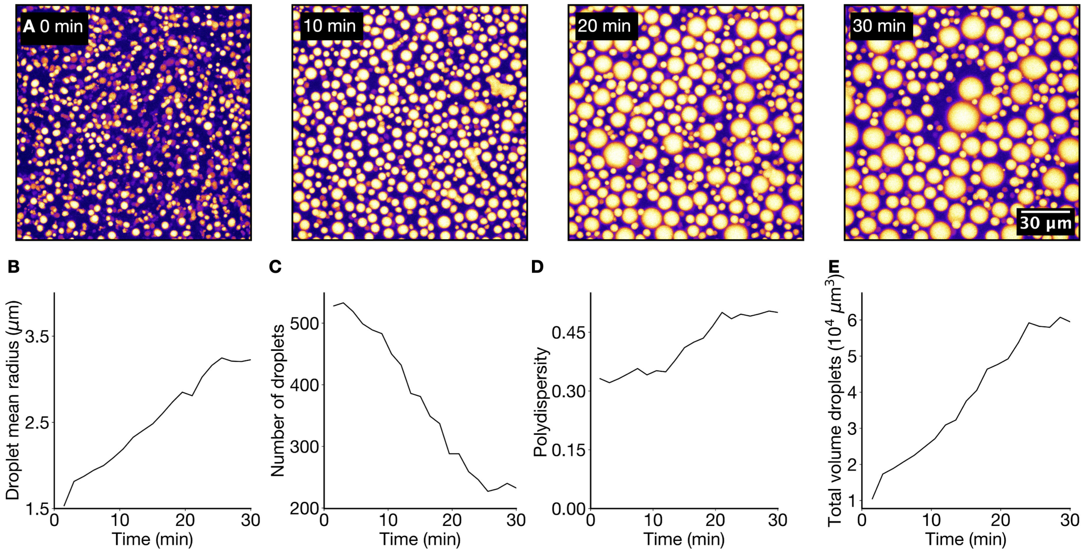

**Supporting Figure 3. Droplets time evolution in steady-state when fusion is not hindered (no gel present).** 15 $\mu$L of aqueous solution with 14 mM peptide, 15 mM PSS and 0.2 $\mu$M sulforhodamine B in MES 200 mM at pH 5.3 were deposited in a well. To start the reaction, 30 $\mu$L of pure DIC were carefully pipetted on top. **A**. Micrographs of active droplets over time when a constant influx of fuel is provided and droplets are free to fuse and sink. Images correspond to z-projections of the 10 $\mu$m imaged to capture the changes in size **B**. Mean droplet radii increases over time as a result of fusion events. **C**. In contrast, the number of droplets decreases. **D**. Droplet size distribution calculated as $\sigma/\mu$ slightly increases over time E. Evolution of total volume of droplets in the imaged volume increases due to droplets sinking.

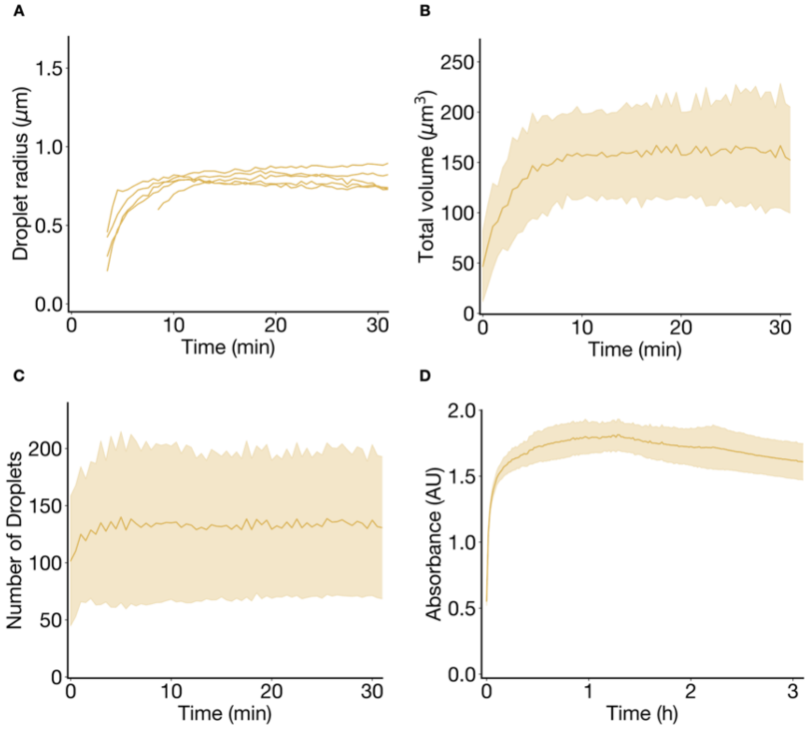

**Supporting Figure 4. Time evolution of active droplets in steady-state with hindered fusion (agarose gel present).** 14 mM peptide, 15 mM polyanion and 0.2 $\mu$M sulforhodamine B were dissolved in 200 mM MES 0.5 w/v $\%$ LMP agarose. A layer of pure DIC was added on top of the gel layer to trigger droplet formation. **A**. Size evolution for each droplet in Figure 2C. Droplets reach the same size independently of the time of nucleation. **B**. The total volume of droplets reaches steady-state after 10 minutes. **C**. The number of droplets in the measured volume rapidly increases and remains constant for the observed time. **D**. Upon droplet formation, the aqueous solution turns turbid. Thus, absorbance at 600 nm was measured to further corroborate constant droplet material on longer timescales. All measurements were performed in triplicates (N=3).

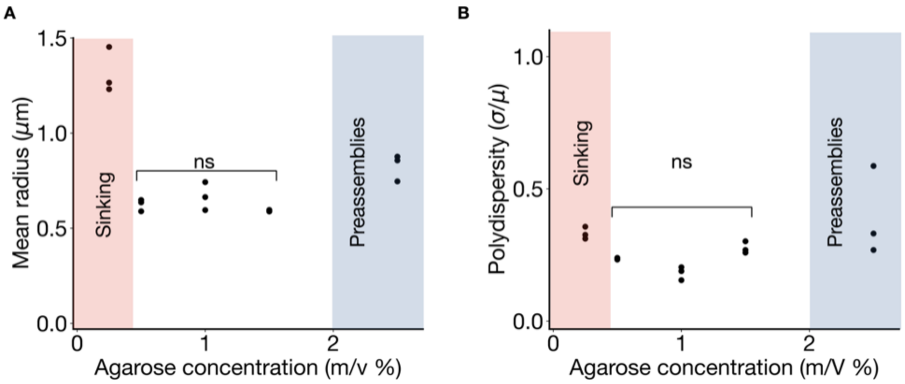

**Supporting Figure 5. Agarose concentration effect on droplet size.** 14 mM peptide, 15 mM polyanion and 0.2 $\mu$M sulforhodamine B were dissolved in 200 mM MES and increasing concentration of LMP agarose (ranging from 0.25 to 2.5 w/v %). A layer of pure DIC was added on top of the gel layer to trigger droplet formation. Concentrations of LMP agarose below 0.5 w/v % were found not sufficient since the droplets were still sinking. Concentrations of LMP agarose above 2 w/v % were causing preassemblies before fuel addition. Concentrations in between showed that the optimal radius of droplets was independent of the agarose concentration. Each point corresponds to an independent sample. Samples were measured in triplicate (N=3). P-values were calculated by comparison to the datapoints at 0.5 w/v %: $p_{1\%}$ = 0.4, 0.06,; $p_{1.5\%}$ = 0.2,0.09 for radius and polydispersity, respectively.

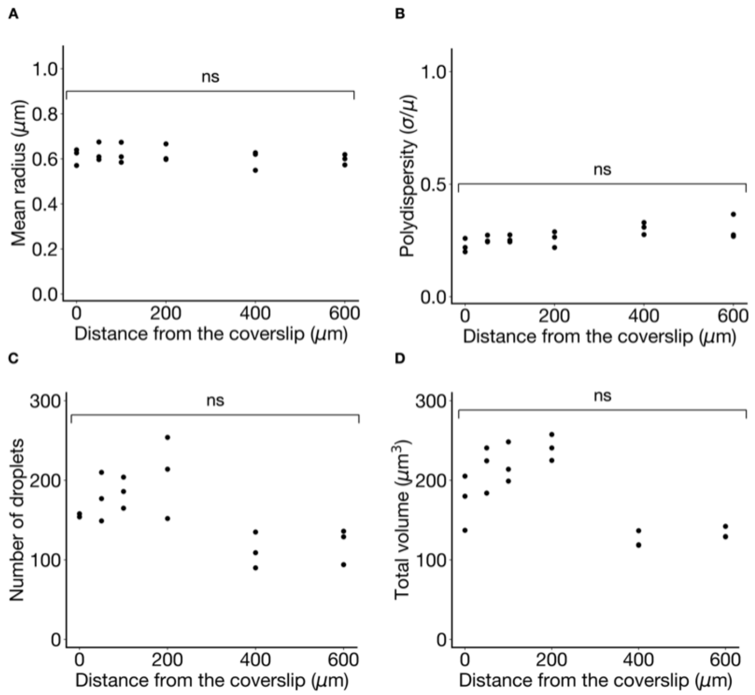

**Supporting Figure 6. Size and distribution of droplets at different heights in the mm-thick gel layer.** To test for fuel gradients, 14 mM peptide, 15 mM polyanion and 0.2 $\mu$M sulforhodamine B were dissolved in 200 mM MES 0.5 w/v % LMP agarose, and layer of pure DIC was added on top of the gel layer to trigger droplet formation. After 1 hour, 10 $\mu$m- thick z-stacks were recorded at different heights. Mean radius of the droplets (**A**), polydispersity (**B**), number of droplets (**C**) and total volume of droplets (**D**) remains constant independently of their distance to the fuel reservoir. Measurements were performed in triplicate (N=3). P-values were calculated by comparison to the datapoints at 0 $\mu$m: $p_{50}$= 0.7, 0.2, 0.2, 0.3; $p_{100}$ = 0.8,0.2,0.1, 0.1; $p_{200}$ = 0.8,0.3,0.06, 0.2; $p_{400}$= 0.7,0.03, 0.1, 0.07;$p_{600}$= 0.6, 0.1, 0.2, 0.1 for radius, polydispersity, total volume and number of droplets, respectively.

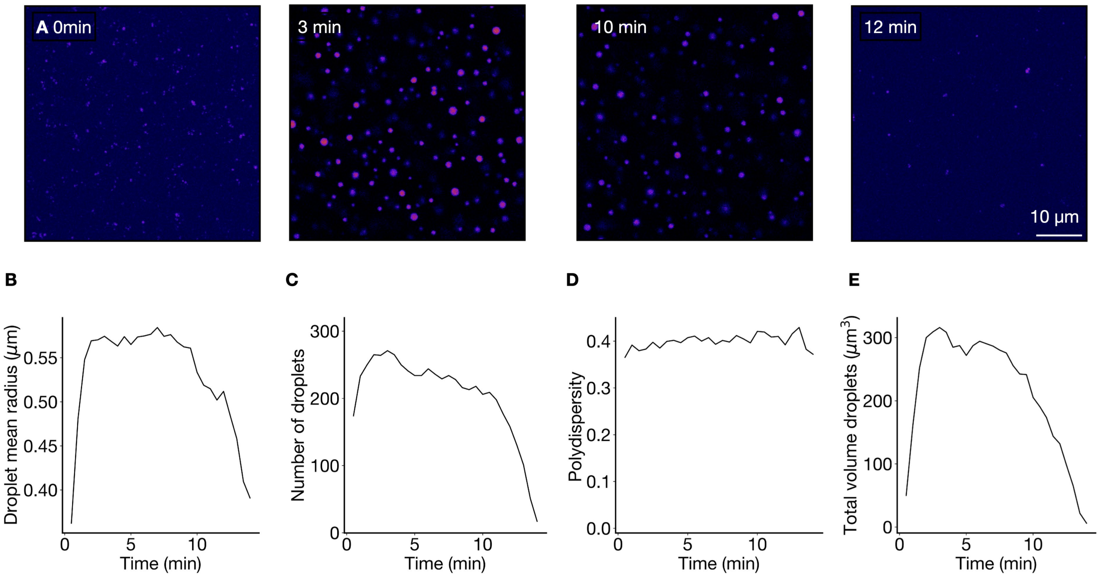

**Supporting Figure 7. Batch fuelled active droplets in agarose gel.** Conditions were 14 mM precursor, 15 mM PSS, and 0.2 $\mu$M sulforhodamine B in 200 mM MES 0.5 w/v % LMP agarose. To trigger coacervation, 25 mM EDC were dropped on top of the gel. After a few minutes, droplets emerged and, once fuel was depleated, they dissolved. **A**. Snapshops of confocal microscopy images showing the emergence, evolution and dissolution of active droplets. **B**. Due to the lack of fusion events, droplets remain relatively small. After 10 minutes, they started dissolving. **C**. Number of droplets rapidly increases upon fuel addition. The number starts decreasing after 10 minutes due to fuel depletion. **D**. Polydispersity remains approximately constant and higher than for droplets sustained in steady-state during the entire experimental time. This suggests droplet size is not controlled by the gel mesh. **E**. Total droplet material follows the same trend as droplet size and number following the emergence of droplets upon fuel addition and their dissolution as fuel is depleted.

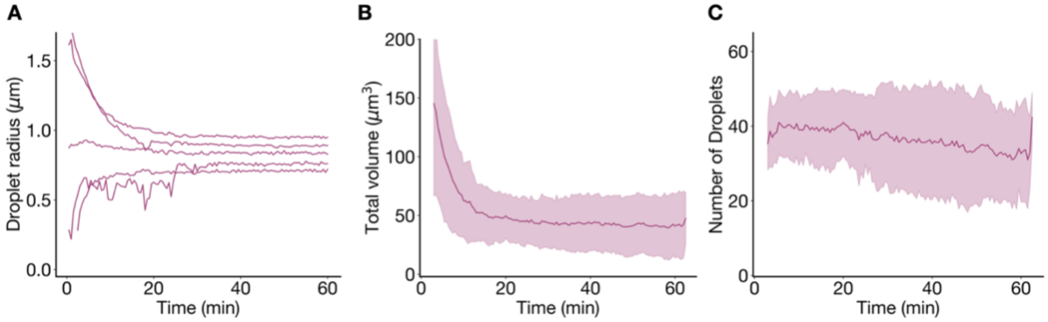

**Supporting Figure 8. Time evolution of prenucleated droplets trapped in agarose gel.** Droplets consisting of 14 mM peptide, 15 mM polyanion and 0.2 $\mu$M sulforhodamine B in 200 mM MES 0.5 w/v % LMP agarose were prenucleated by adding 20 mM EDC while the gel was still liquid. After 3 minutes, a layer of DIC was pipetted on top of the gel to sustain the droplets in steady-state. **A**. Individual droplet's size evolution of droplets shown in Figure 2D. As a result of the prenucleation, the initial size distribution is wider. Over time, small droplets grow while big droplets shrink. **B**. Total droplet material rapidly decreases for the first 20 minutes followed by a stable regime. Such decrease is due to the initial burst in droplet material due to EDC addition while the stabilization indicates that steady-state has been reached. **C**. Number of droplets remains approximately constant during the observed time. All measurements were performed in triplicates (N=3)

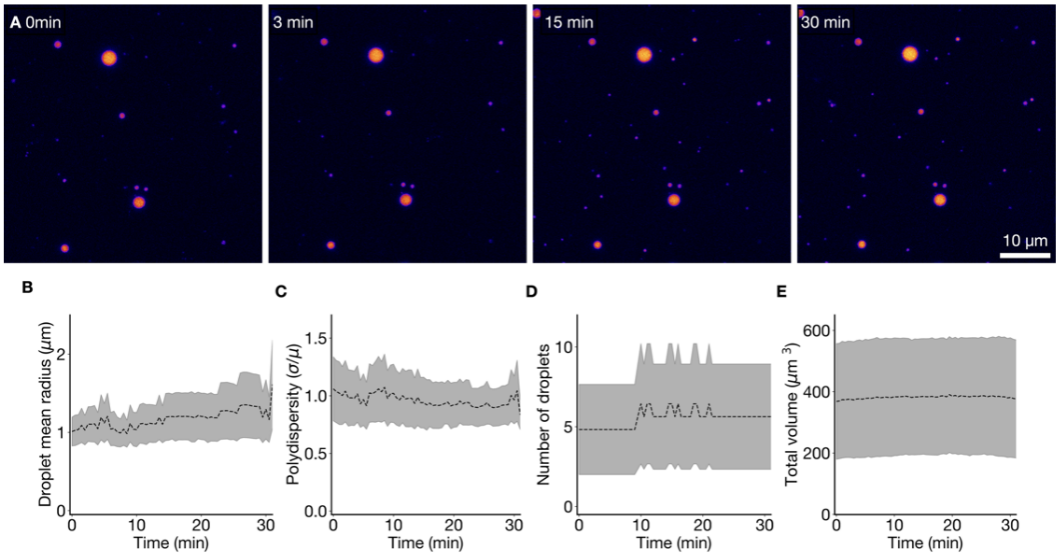

**Supporting Figure 9. Passive droplets evolution over time. A**. Snapshots of the confocal microscopy images showing that droplets' size slowly increases. **B**. Droplet mean radius slightly increases over time, potentially through Oswald ripening. **C**. Polydispersity remains high compared to active droplets, indicating the active nature of the size correction. Number of droplets (**D**) and total volume of droplets (**E**) remain constant over the measured time. Measurements were performed in triplicate (N=3).

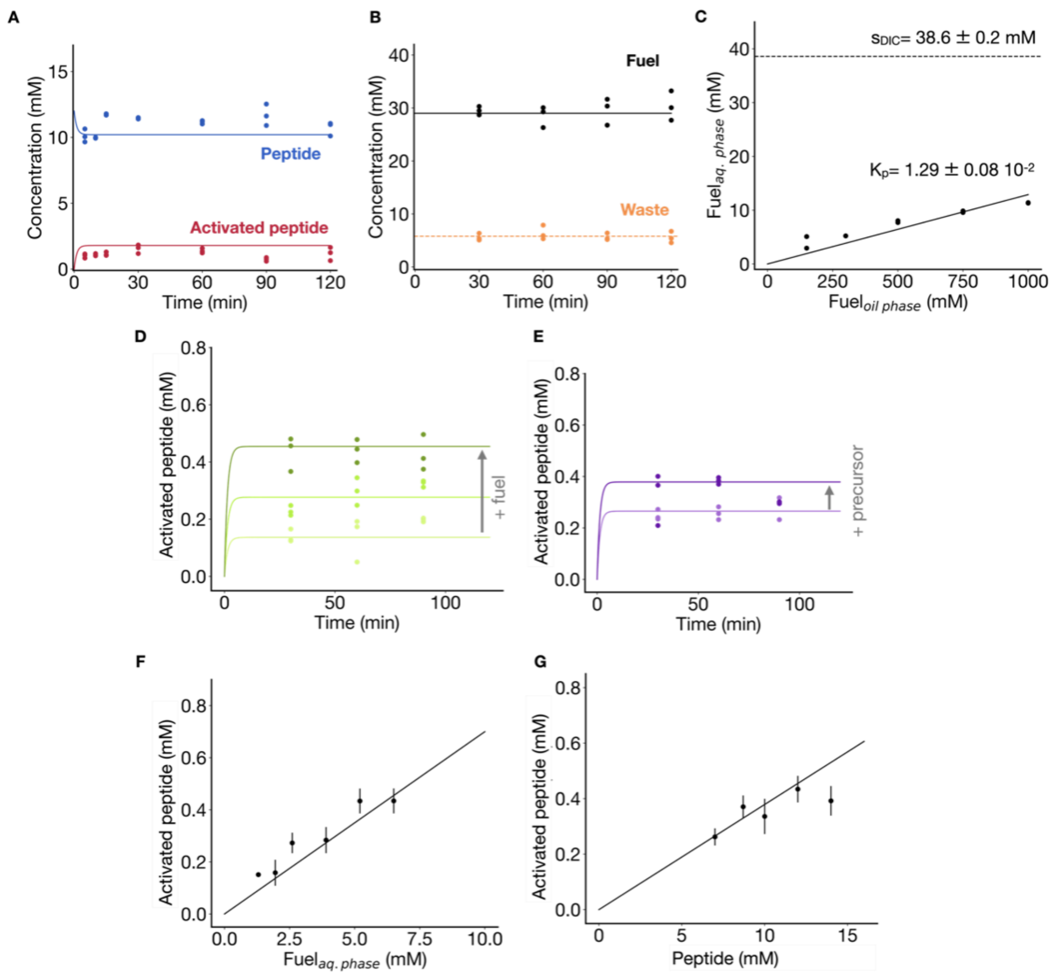

**Supporting Figure 10. Determination of the steady-state kinetics and DIC (fuel) partition coefficient. A.** Concentration of peptide and activated peptide over 2 hours. Conditions were 14 mM peptide in 0.2 M MES at pH 5.3. A layer of DIC was added on top to mimic the experimental setup. Points correspond to measured values determined by HPLC and solid lines to the kinetic model prediction. **B.** Concentration of fuel and waste over 2 hours. Conditions were 14 mM peptide in 0.2 M MES at pH 5.3. A layer of DIC was added on top to mimic the experimental setup. Points correspond to measured values determined by 1H-NMR and solid lines to the kinetic model prediction. In all cases, concentrations stabilized after 15 minutes and the values showed good agreement with the kinetic model. **C.** Determination of fuel concentration in the aqueous phase (0.2 M MES pH 5.3) by 1H-NMR as a function of the concentration in the perfluorinated oil. Data was linearly fitted to obtain Kp. Points correspond to experimental data, the solid line to the Kp fitting, and the dashed line is used as a visual guide of the fuel solubility. Measurements were performed in duplicates (N=2). **D.** Concentration of activated peptide determined by HPLC at different initial fuel concentrations in synthetic cells. The aqueous phase contained 14 mM peptide in 200 mM MES buffer at pH 5.3, and 150 mM (light green), 300 mM (medium green), and 500 mM (dark green) fuel were dissolved in the oil phase. **E.** Concentration of activated peptide determined by HPLC at different initial precursor concentrations in synthetic cells. 8 mM (light purple) and 12 mM (dark purple) precursor were dissolved in the aqueous phase consisting of 0.2 M MES pH 5.3, and the oil phase contained 500 mM of fuel. In all cases, concentrations remained stable for the measured time. Points correspond to measured concentrations and solid lines to the kinetic model prediction, showing good agreement. **F-G.** Stable concentration of activated peptide at different fuel concentrations (**F**) and initial peptide concentration (**G**). Points correspond to the average value measured, bars to the standard deviation, and the solid line to the kinetic model prediction. In both cases, the active peptide concentration increased linearly, and the measured values are in good agreement with the model values. All experiments were performed in triplicate (N=3) except stated otherwise.

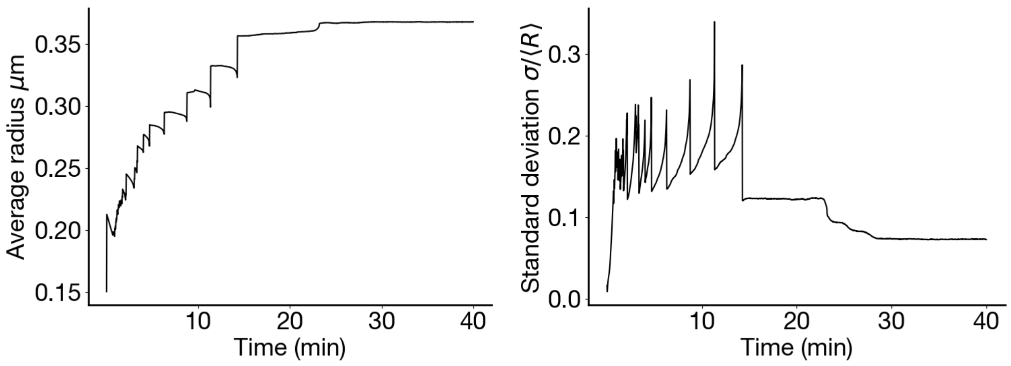

(a) (b)

**Supporting Figure 11. Droplet simulations using the continuous theory described above give a stationary droplet size similar to experiments. (a)** The time evolution of the average droplet radius as observed in phase field simulation of the active emulsion. The saturation in radius with time indicates the steady state. The difference in the saturation radius compared to the experimental value is because the simulations use a two dimensional system compared to the three dimensional experimental system. This affects how the droplet material is distributed among the droplets which is reflected in the average radius. **(b)** The standard deviation with respect to the average droplet radius decays in time, indicating that a nearly mono-disperse state has been achieved.

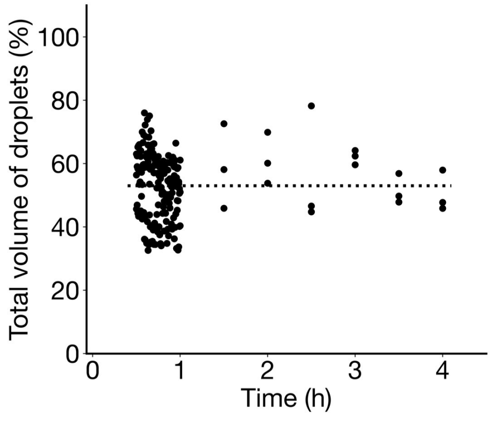

**Supporting Figure 12. Droplet material remains constant when encapsulated in gelified synthetic cells.** Conditions were 14 mM precursor, 15 mM polyanion in 200 mM MES 0.5 w/v % LMP agarose at pH 5.3. 0.2 $\mu$M sulforhodamine B were added to the aqueous phase as droplet dye. The oil phase contained 500 mM DIC as fuel. Total volume was calculated as the percentage of droplet material within a syncell. Each point corresponds to an independent syncell of different size. Measurements were performed in triplicate (N=3).

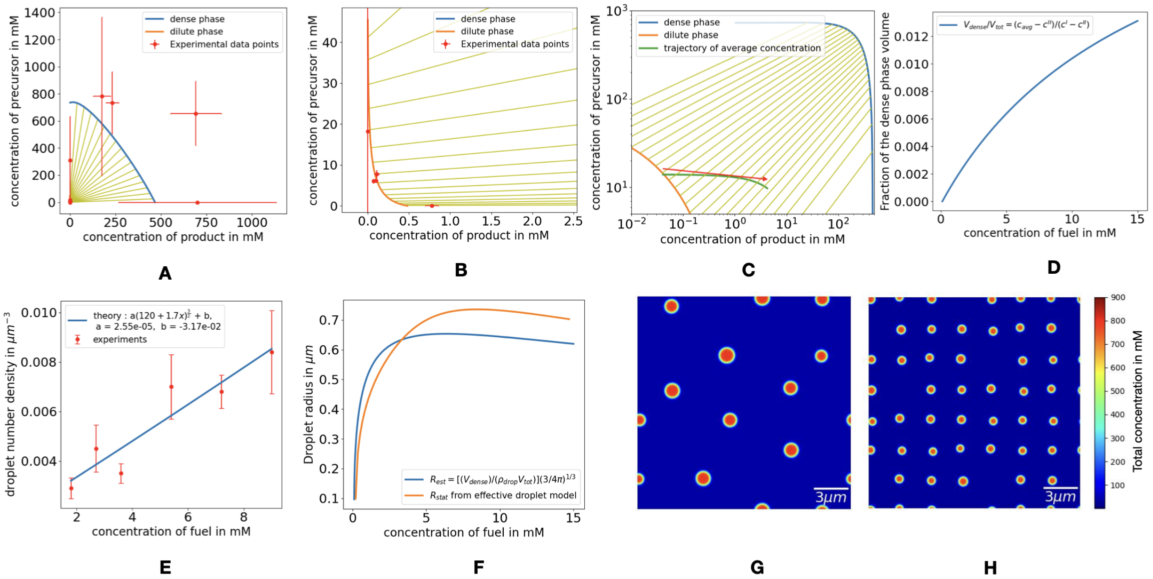

**Supporting Figure 13. : Thermodynamic phase diagram and its role in determining the droplet number density as a function of fuel. (A)** The phase diagram fitted to the experimental concentration values (red dots with arrow bars) from Ref. [1]. The corresponding fitted parameters including the interactions parameters and the molecular volume ratios are given in Table 2. **(B)** Here a zoom in on panel (A) is shows to illustrate the agreement between fit and experimental point along the dilute branch of the phase diagram. **(C)** The trajectory of the average concentration as we vary the fuel from 0.1mM to 15mM is depicted in green. The red arrow represents the direction of increasing fuel. As we increase fuel, the distance of the average concentration from the dilute binodal (orange) increases, leading to an increase in the fractional volume of dense phase. **(D)** Fraction of the dense phase volume $V_{\text{dense}}/V_{\text{tot}} = (c_{\text{avg}} - c^{\text{II}})/(c^{\text{I}} - c^{\text{II}})$ plotted corresponding to the green trajectory in (**C**). **(E)** The droplet number density expressed as a function of fuel concentration as per Eq.((25)). The factor 'a' has been defined in Eq.(25). The behaviour of the droplet number density for very low fuel values may no longer be described by the relation in Eq.(25) since droplets disappear altogether below a threshold fuel concentration. So we use an additional parameter 'b' which can account for this effect. **(F)** The figure compares the stationary radius $R_{\text{stat}}$ from the effective droplet model with the estimate of the stationary radius $R_{\text{est}}$ from the phase diagram and the droplet number density shown in panels **(C)**,**(D)** and **(E)** respectively. The agreement between the curves confirms to a large extent that the condensed volume and chemical activity set the stationary droplet radius as per Eq. (22). The slight discrepancy between the two curves can be attributed to the presence of concentration gradients in the active system (orange curve), which are implicitly assumed to be absent while estimating the radius using the phase diagram (blue curve). Images in **(G)** and **(H)** are snapshots of the phase field simulations for fuel concentrations of 8mM **(G)** and 19mM **(H)** at steady state. It is observed that the droplet number density is higher for a higher fuel concentration as expected through theoretical predictions.

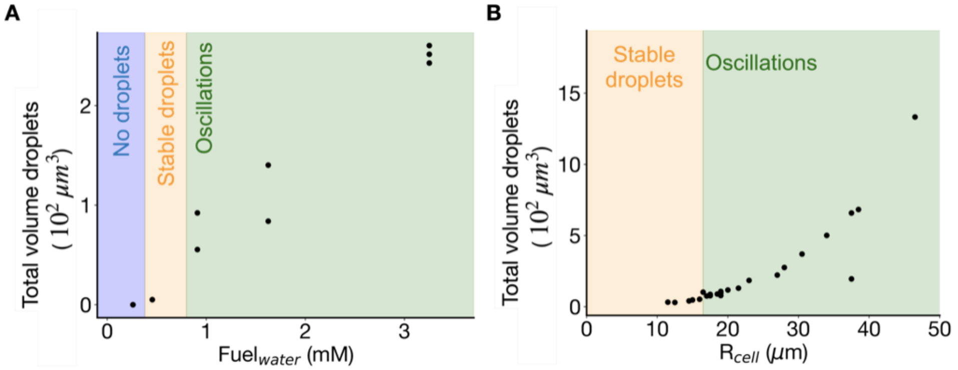

**Supporting Figure 14. Total volume of droplets in synthetic cells at varying conditions. A.** Total volume of droplets increases with the concentration of fuel due to an increase in the steady-state level of activated peptide. **B.** The total droplet material also increases with the size of the synthetic cell.

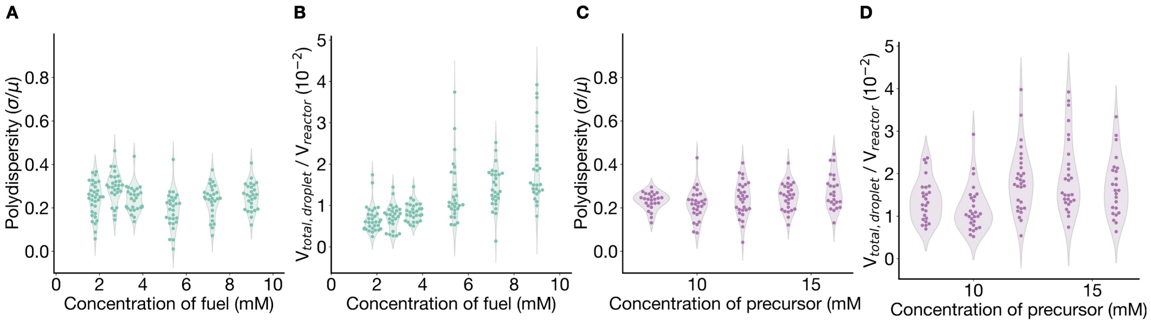

**Supporting Figure 15. Polydispersity and droplet volume at different steady-states in synthetic cells.** Polydispersity at different fuel (**A**) and precursor (**C**) concentrations remains constant. The amount of droplet material slightly increases as fuel (**B**) and precursor (**D**) concentrations are increased. Each point corresponds to a synthetic cell of a different size from three independent samples.

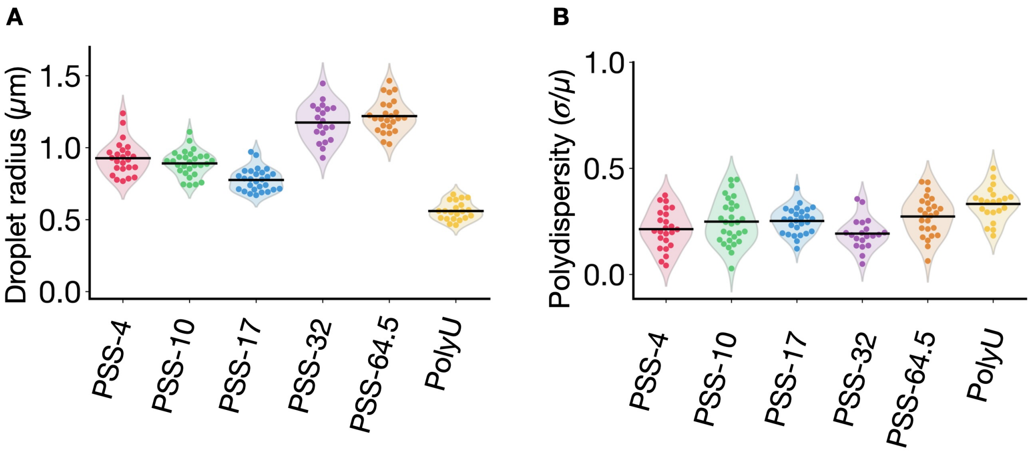

**Supporting Figure 16. Influence of polyanion.** 14 mM precursor, 15 mM polyanion and 0.2 $\mu$M sulforhodamine were dissolved in 0.5 w/v % LMP agarose at pH 5.3 200 mM MES. The oil phase contained 500 mM DIC as fuel and synthetic cells of different sizes were produced. **A**. Droplet radius depends on the length and chemical structure of the polyanion, probably due to the different interactions with the activated peptide. In contrast, polydispersity (**B**) remains similar, suggesting that the size control mechanism is taking place in all samples. Each point corresponds to a synthetic cell of a different size from three independent samples.
